# Brainstem sensing of multiple body signals during food consumption

**DOI:** 10.1101/2025.04.28.651046

**Authors:** Rachel A. Essner, Kiersten Ruda, Hannah J. Choh, Hakan Kucukdereli, Oren Amsalem, Julia Edelhaus, Sverre Grødem, Kristian K. Lensjø, Teresa E. Lever, Mark L. Andermann

**Author notes:** These authors contributed equally (Rachel A. Essner, Kiersten Ruda). Correspondence (M.L.A.).

## Abstract

Studies of body-to-brain communication often examine one stimulus or organ at a time, yet the brain must integrate many body signals during behavior. For example, food consumption generates diverse oral and post-oral chemical and mechanical signals transduced by well-characterized peripheral neuronal pathways. Far less is known about how these and other bodily signals are integrated and organized in the brainstem lateral parabrachial nucleus (LPBN), a key interoceptive sensory hub. We established methods to image the activity of 1000s of neurons throughout a large region of mouse LPBN. Food consumption drove a seconds-long wave of activity across LPBN, with dynamics mirroring the movement of food through the upper gastrointestinal tract observed using X-ray fluoroscopy. By imaging the same neurons across days, we found that spatially clustered subsets of neurons encoded oral signals, stomach filling, visceral malaise, arousal, and/or body movement. Moreover, only certain subsets were modulated by cortical input. Together, these experiments reveal a functional specialization in the LPBN that integrates contextual information from the body to guide behavior.

## Introduction

Interoception, the sensing of internal body signals, is critical for regulating motivated behaviors, such as food consumption, to maintain physiological homeostasis and ensure survival. Effective regulation of food intake by the brain requires tracking and integrating diverse body signals from multiple organs within the gastrointestinal (GI) tract. For example, the decision to continue to eat is informed by chemical signaling from tastants in the mouth and nutrient absorption in the intestines, and by mechanical signaling from esophageal, stomach, and intestinal stretch. Hormones, including cholecystokinin (CCK), glucagon-like peptide-1 (GLP1), and amylin, are also released from various organs in response to a meal^1^. How these many peripheral signals are integrated by the brain during natural behaviors remains largely unknown.

The brainstem lateral parabrachial nucleus (LPBN) is a key hub that receives sensory information about the body from vagal (via the nucleus of the solitary tract, NTS), spinal (via the spinoparabrachial tract), and humoral (via the area postrema, AP) pathways. The LPBN integrates these sensory signals and drives relevant changes in behavior through bidirectional communication with forebrain regions such as the hypothalamus, amygdala, and cortex^2–4^. As such, the LPBN is involved in the regulation of various interoceptive processes, including appetite^5–7^, body temperature^8,9^, fluid balance^10^, breathing^11–13^, pain^4,14–18^, and itch^19,20^. Previous work has characterized multiple genetically defined cell types that are mostly restricted to different subregions of the LPBN. These genetic subpopulations also exhibit projections to distinct downstream brain areas and mediate different functions of LPBN^21–23^. For example, calcitonin gene-related peptide (CGRP)-expressing neurons reside within the external lateral region (PBNel) and have been coined “alarm” neurons as they mediate behavioral responses to various bodily threats, such as visceral illness, pain, and hypercapnia^3^. Other well-characterized examples involve prodynorphin (Pdyn)-expressing neurons in the dorsal lateral region (PBNdl), which mediate responses to upper GI tract stretch and body heating^8,24,25^, and Satb2-expressing neurons in the ventral-lateral region that encode taste^26,27^.

Despite the progress in genetic characterization of LPBN neuron subtypes, it is unlikely that any one of these genetically defined populations tracks only a single type of body signal. Rather, multiorgan and multimodal signals are likely integrated and transformed in LPBN to inform the brain about distinct bodily states. To build a more holistic view of this sensory coding in LPBN requires understanding neural sensitivities to the body’s sensory landscape during various behaviors and to more conventional, experimentally controlled stimuli (e.g., anesthetized sensory mapping^28^). In the case of food intake, this requires tracking the dynamics of food through the GI tract during consumption, as well as other body and state-related changes associated with feeding, such as an increase in arousal and a decrease in locomotion – both of which induce activity in LPBN neurons in a manner distinct from GI tract chemo- and mechano-sensation^11,13,29,30^. Finally, in contrast to external sensory stimuli, internal stimuli associated with body states (e.g., satiation, malaise) are not rapidly reversible and cannot be presented in sequence in the same neural recording session. Therefore, we require additional innovations to comprehensively profile interoceptive sensory properties via longitudinal recordings of LPBN neurons across sessions.

To address these challenges, we first assessed GI tract dynamics during food consumption using X-ray videofluoroscopy in mice. We found that liquid food moves with relatively stereotyped timing from the mouth to the stomach and proximal small intestine in seconds, and it also accumulates in the stomach across many minutes. Separately, we imaged the activity of 1000s of LPBN neurons in response to several feeding related, malaise-inducing, and painful stimuli across days while tracking behavioral variables including licking, locomotion, and pupil area. This approach, coupled with subsequent anesthetized mapping of stomach stretch, revealed spatially organized groups of neurons that respond to distinct aspects of food intake, visceral malaise, and locomotion. More generally, our work highlights a novel approach to examine how neurons in the LPBN and elsewhere organize and integrate multiple body signals in awake animals.

## Results

### Gastrointestinal tract dynamics during feeding

To better understand how the brain responds to interoceptive signals during feeding, we first sought to define the dynamics with which food moves through the upper GI tract. To this end, we developed a protocol for X-ray videofluoroscopy^31^ in head-fixed awake mice to observe the ingestion of food through the GI tract using the same feeding protocol that we then employed for neural recordings below. Mice (n=4) received small drops (20 μl) of Vanilla Ensure every 20 s **(Figure 1A).** While bone tissue is readily apparent under video fluoroscopy (**Figure 1B),** mixing Ensure with the radiopaque contrast agent iohexol allowed us to visualize the movement and accumulation of Ensure from the mouth and throat to the stomach and proximal small intestine **(Figure 1F and Videos S1-2).**

**Figure 1.**
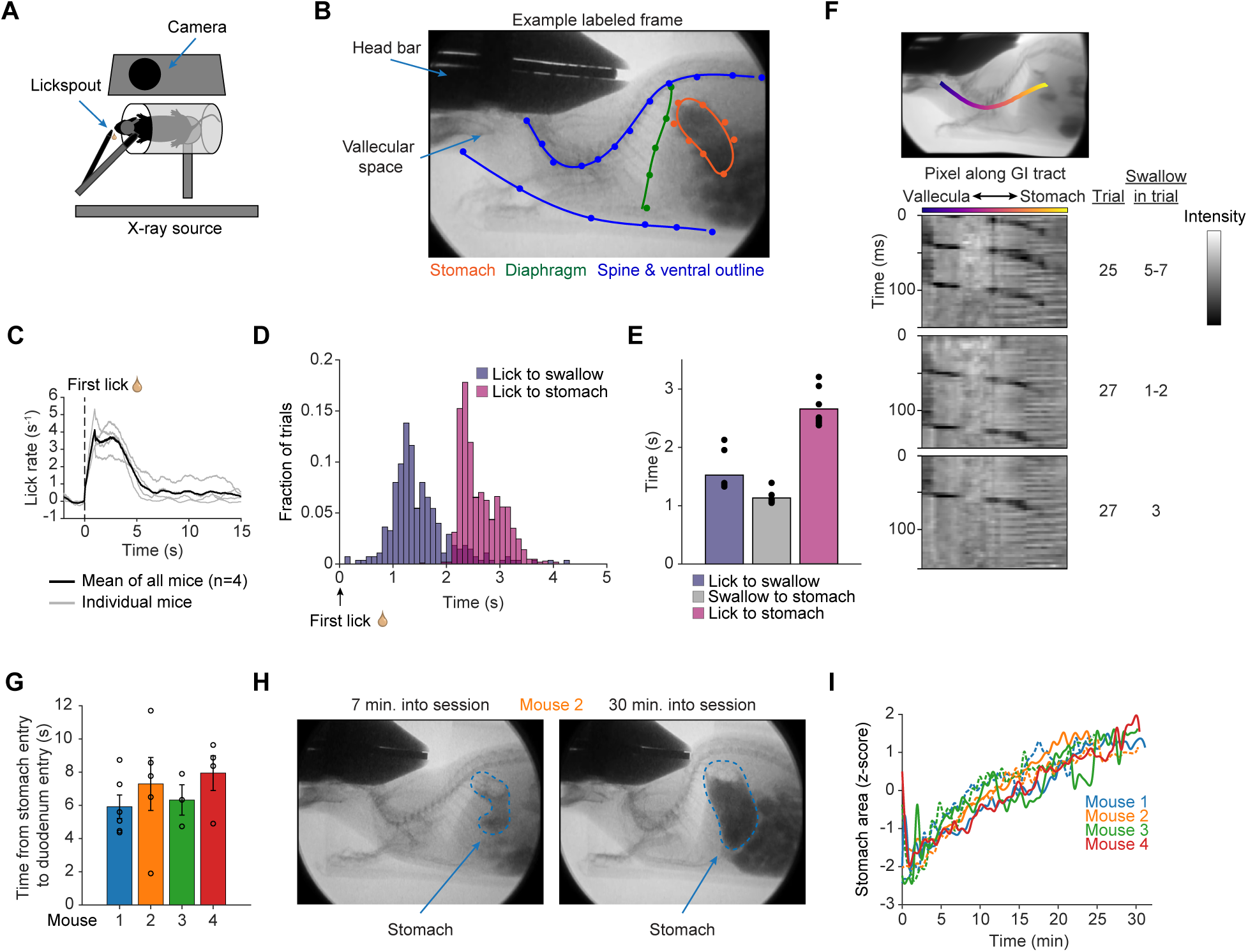
Gastrointestinal tract dynamics during feeding. (A) Bird’s eye view of head-fixed setup for food (Ensure) intake and lick recording during X-ray fluoroscopy. (B) Side-on X-ray image from an example mouse showing DeepLabCut labeled points. (C) Mean lick rate across all trials from all mice (40-52 trials/mouse from 4 mice) aligned to the first lick of every trial. Black: Mean across all mice; Gray: Mean across all trials from an individual mouse. (D) Distribution of time from first lick to first swallow and from first lick to first stomach entry of a food bolus, for all trials across all mice (first lick to swallow: n = 275 trials from 4 mice; lick to stomach entry: n = 1056 trials). See also **Supplemental Video S1.** (E) Quantification of the mean time from first lick to first swallow, swallow to stomach, and first lick to stomach across mice (n = 4). Each dot is the mean of 11-52 trials for first lick to swallow or 37-227 trials for swallow to stomach entry from a single mouse. (F) Top: X-ray image with the upper GI tract labeled in color to highlight the path of an Ensure bolus from the mouth to stomach. Bottom: Kymograph showing the movement of a bolus of Ensure from mouth to stomach (dark band moving down and to the right) for example swallows from an example mouse. Swallows are taken from 3 separate trials as noted. (G) Quantification of the average time between the first entry of an Ensure bolus into the stomach and the first duodenum entry (n = 3-6 trials for each mouse). Error bars are SD. See also **Supplemental Video S2**. (H) X-ray images showing the change in stomach volume (and filling of the intestines) from the beginning (left) to the end (right) of a recording session in an example mouse. (I) Traces showing the increase in stomach area (calculated from DLC-labeled points; see Methods) over time on two separate recording days for each mouse. Mice ingested ∼1.5 ml of Ensure per session (see **Figure S1D** for absolute area). Solid line: Day 1; dashed line: Day 2.

We observed gastrointestinal dynamics across timescales from seconds to many minutes. Upon Ensure delivery, the food-restricted mice readily licked for Ensure for ∼4 s **(Figure 1C).** Ensure accumulated in the vallecular space at the back of the mouth during licking **(Figure 1B)**, and the first bolus of Ensure was swallowed, on average, 1.52 +/- 0.32 s (mean +/- SEM) after the first lick of a trial **(Figure 1D-E).** We also observed multiple swallows per trial (3.67 +/- 0.43 on average, **Figure S1A)**. The average esophageal transit time was 1.15 +/- 0.11 s, resulting in a mean transit time from licking onset to stomach entry of 2.68 +/- 0.30 s **(Figure 1E-F).** These ingestion kinetics were largely consistent across recording days from the same mouse and across mice (**Figure S1B-C)**.

We next sought to estimate the movement of Ensure into and out of the stomach, as previous studies have only investigated stomach emptying on the timescale of minutes^32,33^. We analyzed a subset of the first 7 trials in each session, which had higher contrast for observing movement through the stomach due to a lack of accumulated iohexol. We observed clear emptying of food from the stomach into the proximal small intestine (duodenum, **Video S2**), which senses nutrients and rapidly influences the brain’s reward pathways^34–37^. For those trials, we quantified the time between the first bolus entry into the stomach and the first instance of movement into the duodenum **(Figure 1G).** These data indicate a surprisingly fast movement of a portion of the ingested liquid food out of the stomach, with a delay from entry into the stomach to arrival in the duodenum in as little as 5.92 +/- 0.71 s (mean +/- SEM for other mice: 7.29 +/- 1.6 s, 6.32 +/- 0.91 s, and 7.94 +/- 1.0 s). Inspection of movies from single trials indicated that some of the ingested fluid is routed to the forestomach for temporary storage **(Videos S2-3)**, even on early trials within a session, and that emptying of food into the duodenum does not occur on every trial (data not shown). Overall, these analyses show that liquid food can take as little as ∼9-11 s to travel through the upper GI tract from initial licking to arrival in the duodenum, where post-ingestive sensing of nutrients occurs^34–37^.

On a slower timescale, we observed dramatic gastric expansion over the course of each 30-minute recording session (∼1-1.5 mL Ensure consumption; **Figure 1H-I and Video S3**). To quantify this change in stomach volume, we used DeepLabCut^38^ to track points along the perimeter of the stomach and estimated stomach area using a spline fit of these points (**Figure 1B and Video S4**). Stomach area expanded as much as 4.9 times during the feeding paradigm **(Figure S1D)**, consistent with a previous report of stomach volume changes when mice consume chow or liquid following fasting^24^. This expansion highlights the substantial changes in stomach size (and associated accommodation of the forestomach) during food consumption. Together, these results help define the natural events that occur across seconds to minutes during consumption.

### LPBN neurons respond to ingestion with varied temporal dynamics

We wondered whether different LPBN neurons track the arrival of food in different regions of the GI tract, and how this relates to each neuron’s sensitivity to other stimuli and behaviors. To this end, we developed a novel preparation to record activity from thousands of LPBN neurons using chronic two-photon calcium imaging. In contrast to prior studies focusing on sparse genetic subtypes^13,20,24,27,39^, we sought a broad characterization of neurons across a large region of LPBN. Initial experiments showed strong background fluorescence (i.e., neuropil contamination) in LPBN when using cytosolic expression of the GCaMP6s calcium indicator. Thus, we expressed soma-restricted GCaMP8s (ribosome-tethered, ribo-GCaMP8s^40,41^) in all LPBN neurons and implanted a 1-mm diameter gradient index (GRIN) lens at an angle just above the surface of LPBN to avoid damage to the LPBN itself **(Figure 2A-B).** This approach allowed us to image from a large portion of LPBN (825 µm x 530 µm per FOV on avg.), including the PBNel and PBNdl subregions (which contain CGRP and Pdyn neurons, respectively). We confirmed the imaging locations using 3D histological reconstruction of the GRIN lens tract in coronal sections followed by alignment to the Allen Mouse Brain Common Coordinate Framework **(Figures 2C and S2A-B)**. Overall, we recorded the activity of approximately 3600 neurons across two depths within LPBN from 9 mice (∼400 neurons/recording).

**Figure 2.**
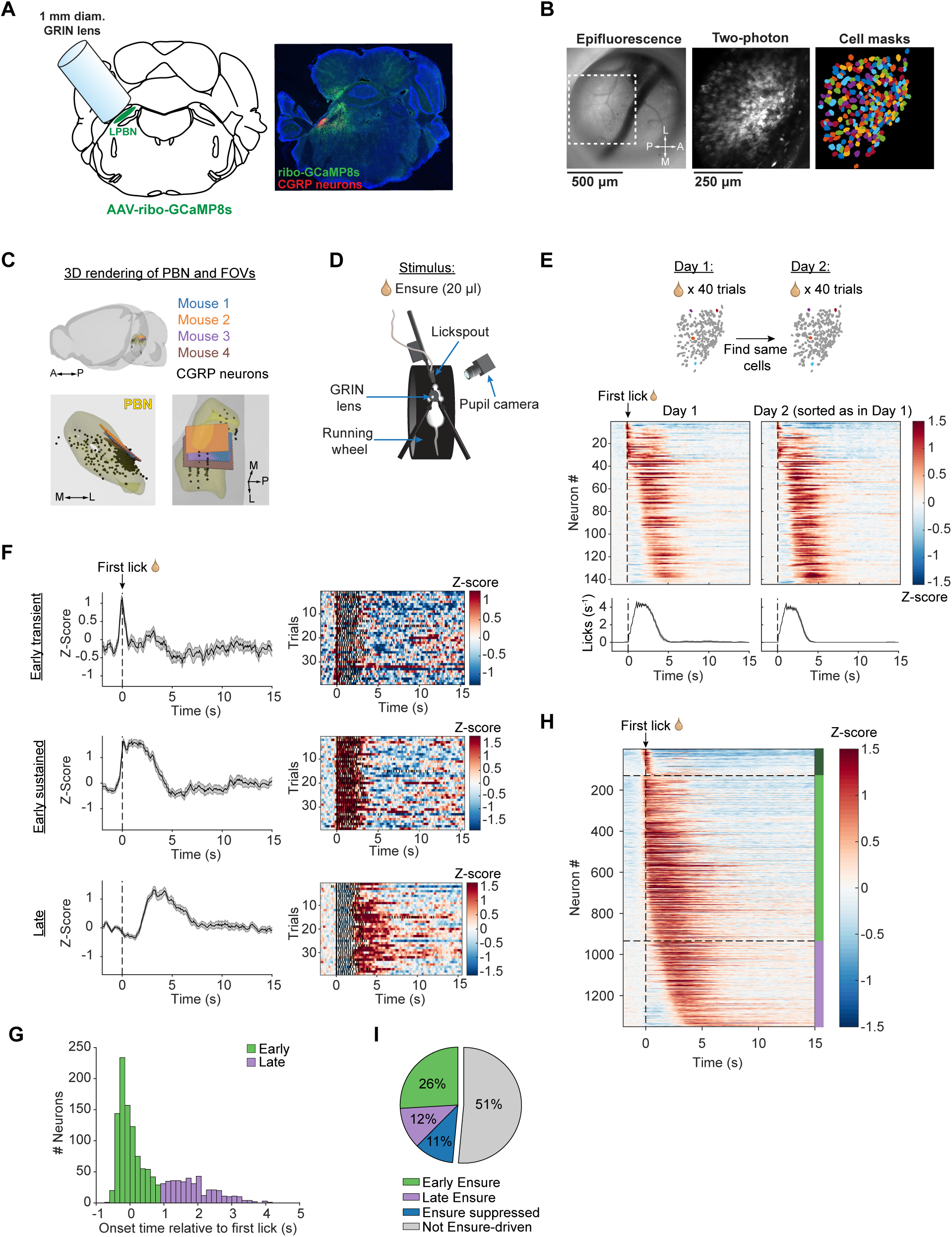
LPBN neurons respond to ingestion with varied temporal dynamics. (A) Two-photon calcium imaging of LPBN through a GRIN lens. Left: Schematic of ribo-GCaMP8s expression and 1-mm diameter GRIN lens implantation at an angle above LPBN. Right: Example coronal section showing GRIN lens tract above ribo-GCaMP8s-expressing LPBN neurons (green). PBNel CGRP neurons (red) are labeled for reference. (B) Example imaging field of view. Left: Epifluorescence image of LPBN surface and vasculature through a GRIN lens. P, posterior; A, anterior; M, medial; L, lateral. Middle: Two-photon image of boxed region in left image. Right: Cell masks extracted from middle image using Cellpose^109^. (C) 3D rendering of fields of view (FOVs) from 4 example mice after alignment to the Allen Mouse Brain Common Coordinate Framework. Gray: 3D outline of mouse brain. Yellow: 3D outline of entire PBN. Colored rectangles: FOVs from 4 example mice. Black: CGRP-expressing neurons to highlight location of PBNel. A, anterior; P, posterior; M, medial; L, lateral. (D) Head-fixed setup for food (Ensure) intake during two-photon calcium imaging while recording locomotion, licking, and pupil size. (E) Top: Schematic for analyzing neural responses across two consecutive days of imaging. Middle: Heatmaps showing the mean response (n=40 trials) of single Ensure-activated neurons (n=144) from an example mouse. Each row is the timecourse of an individual neuron. Activity is aligned to the first lick of each trial, which occurs at T=0 s (dashed line), and normalized to the 2 s baseline period prior to the first lick. Middle left: Heatmap of responses from the imaging session on Day 1. Middle right: Heatmap of responses of the same neurons from the imaging session on the following day, Day 2. Neuron order is sorted by onset time (see Methods) on Day 1. Bottom: Mean lick traces (n=40 trials) for Days 1 and 2, aligned to the first lick of each trial. (F) Mean Z-scored activity traces (left) and single-trial activity heatmaps (right) for example Early transient, Early sustained, and Late onset Ensure-responsive neurons. Black tick marks in heatmaps indicate individual licks. (G) Histogram showing the distribution of onset times of all Ensure-activated neurons (n=1351 neurons from 9 mice). An onset time of 0.9 s divides the neurons into Early (green) and Late (purple) groups. See also **Figure S2E**. (H) Heatmap showing the mean response of all Ensure-activated neurons (n=1351 neurons from 9 mice), aligned to the first lick and sorted by onset time, as in (E). Color bar to right indicates Early transient (dark green), Early sustained (green), and Late (purple) neuron groups. (I) Pie chart demonstrating the percentage of each neuron type in the whole population of recorded LPBN neurons (n = 3602 neurons from 9 mice). For all panels, data is displayed as mean +/-SEM.

To determine how LPBN neurons respond during liquid food (Ensure) consumption, we imaged LPBN during a similar feeding paradigm to that in **Figure 1**. Mice were head-fixed atop a running wheel, and we recorded licking, locomotion, and pupil area (a measure of arousal^42,43^) during imaging sessions **(Figures 2D and S2H-J)**. We found that Ensure consumption drove a wave of activity in LPBN, where some activated neurons responded at the onset of licking while others only began to respond 1-4 s later **(Figure 2E-H and Video S5).** These temporal dynamics were present in every field of view (FOV) across all nine imaged mice **(Figure S2C)**, and we confirmed that these FOVs were near each other in LPBN using the Allen Brain Mouse Atlas mapping described above **(Figures 2C and S2A-B)**. Further, the distinct response dynamics of each neuron were consistent across feeding trials within a session, and across two consecutive daily sessions (in instances where we could track the same neurons across days; **Figures 2E-F and S2D-E).** Overall, Ensure consumption significantly activated 38% (1352/3602) of all imaged LPBN neurons (**Figure 2I**; across n = 9 mice; see Methods). Feeding also suppressed activity in a smaller subset of neurons, and this suppression exhibited variable dynamics across neurons (**Figure S2G**). However, since suppressed neurons were considerably less common (11%; 397/3602, **Figure 2I**), we focused on activated neurons in subsequent analyses (note 51% of imaged LPBN neurons were not driven by Ensure, **Figure 2I**).

We divided the feeding-activated neurons into three groups based on response characteristics: those with (1) Early onset and transient responses (onset < 0.9 s after first lick, duration ≤ 0.9 s), (2) Early onset and sustained responses (onset < 0.9 s, duration > 0.9 s), and (3) Late onset (onset ≥ 0.9 s, “Late Ensure” neurons) **(Figures 2F-I and S2F**; see Methods). Given the small number of early and transient neurons relative to the entire population (9.6%, 130/1351 Ensure-activated neurons), we grouped these neurons with the early and sustained neurons for subsequent analyses (together, “Early Ensure” neurons). All Early Ensure neurons showed response onsets that coincide with or precede licking; thus, these neurons may encode motor signals associated with licking, taste, mechanosensory signals from the mouth/tongue, and/or food-anticipatory signals. In contrast, Late Ensure neurons respond to food during or after the first swallow of each bout (76% of swallows occurred from 0.9-1.9 s after the first lick, see **Figures 1D-E)**, suggesting that these neurons respond to mechanosensory signals from the esophagus and/or stomach, and/or predict upcoming stimuli in these regions (see Discussion). In fact, many Late Ensure neurons begin to respond 2-4 s after the first lick, indicating that these neurons may be sensitive to entry of food into the stomach (mean time from first lick to stomach entry: ∼2.7 s, compare **Figures 1D-E and 2G-H).**

Thus, our large-scale imaging approach reveals a functional specialization of LPBN neurons that respond during and immediately following the moment of consumption, which we hypothesize is due to distinct sensitivities to various sensory and motor signals from oral and post-ingestive aspects of feeding. To test this hypothesis, we directly assessed the sensitivity of these same recorded LPBN neurons to post-ingestive GI signals, as described below.

### LPBN neurons respond to post-ingestive signals

First, we delivered intraperitoneal (i.p.) injections of LiCl, a compound that induces visceral malaise. In addition, during a terminal anesthetized experiment, we implanted a balloon in the stomach to examine responses to stomach stretch. Both LiCl and stomach stretch have been shown to drive activity in LPBN^6,24,35,44^, yet it is unclear whether the same or different neurons respond to these two stimuli, and whether responsive neurons are also activated by Ensure consumption with specific dynamics. Since we propose that Late Ensure neurons are sensitive to mechanosensory signals from the stomach, we hypothesized that they would be more responsive to stomach stretch than Early Ensure neurons or those not driven by Ensure.

We first recorded responses to Ensure consumption as in **Figure 2**, then delivered a control i.p. injection of 0.9% saline, and finally an i.p. injection of LiCl (0.2M, 14 ml/kg; **Figure 3A**, see Methods). Many neurons (28%, 1235/4447) showed robust increases in GCaMP8s fluorescence beginning 3-5 minutes after LiCl injection and persisting for at least 25 minutes **(Figure 3B-C)**. These calcium dynamics are consistent with previous studies showing increased Fos expression in LPBN after LiCl injection^6,35,44,45^. We also observed a smaller fraction of neurons that decreased fluorescence in response to LiCl (18%, 790/4447, **Figure 3C)**. In contrast, LiCl-activated neurons showed much less activation in the 10 minutes following saline injection, indicating that LiCl responses are not due to injection-induced pain, stress, or motion artifacts **(Figures 3D and S3A).** Next, we compared these LiCl responses to those during Ensure consumption. We found that Late Ensure neurons tended to exhibit stronger activation by LiCl than Early Ensure neurons **(Figures 3E-F and S3B).** Further, even though only 28.4% of the neurons responsive to Ensure were Late Ensure neurons, 54.6% of neurons responsive to both Ensure and LiCl were late Ensure neurons **(Figure 3G).**

**Figure 3.**
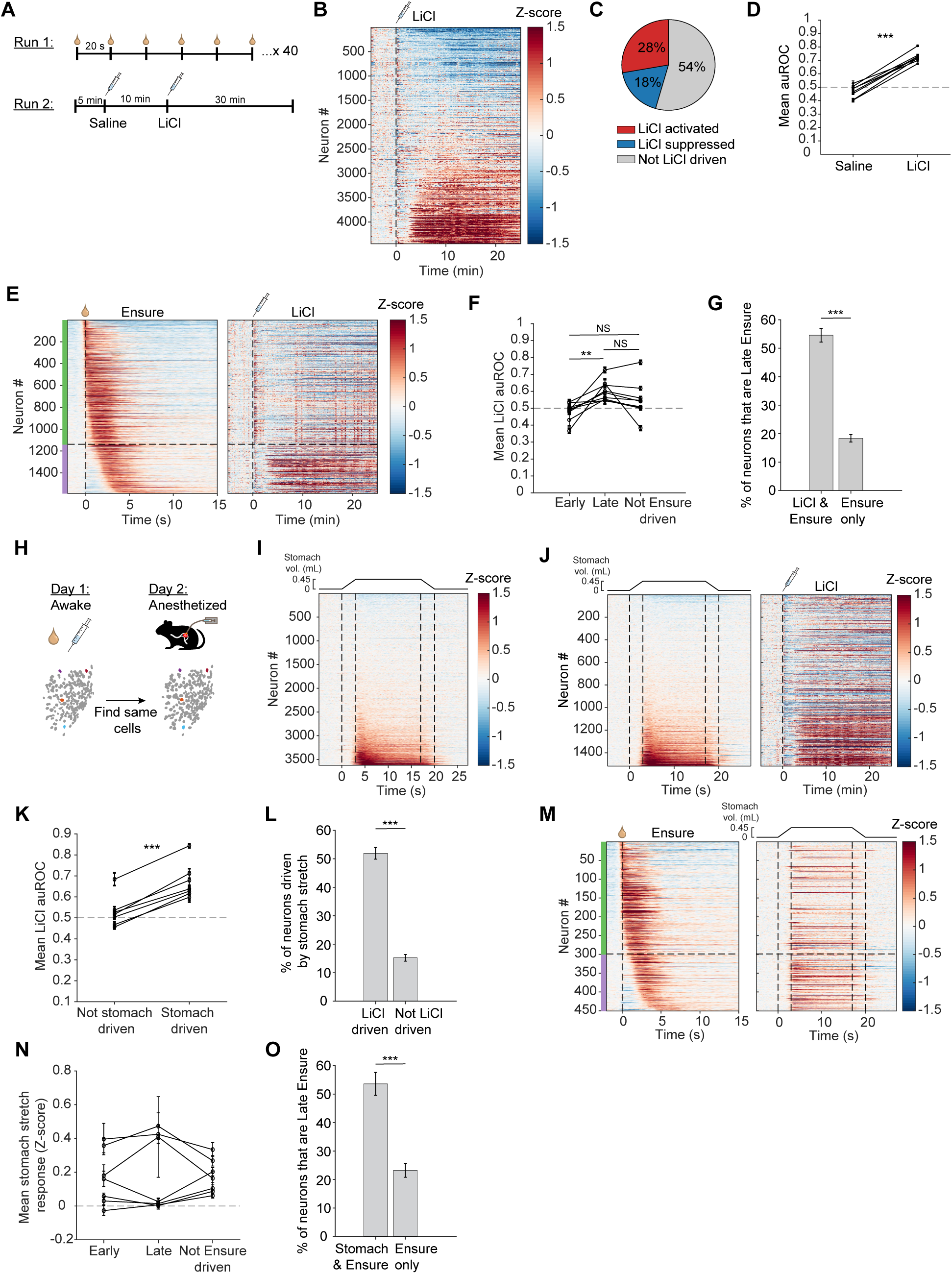
LPBN neurons respond to post-ingestive signals. (A) Schematic of experiment for analyzing LPBN responses to i.p. drug administration. (B) Heatmap showing the response of single LPBN neurons (n = 4447 neurons from 9 mice) to LiCl injection. Each row is the response of a single neuron, and neurons are sorted by auROC value from the baseline activity in the 5 min prior to injection and the activity in the 25 min post injection. Injection occurred at time T=0 min. (C) Pie chart demonstrating the percentage of LiCl activated, LiCl suppressed, and not LiCl driven neurons within the entire population of recorded neurons. LiCl activated: 1235/4447; LiCl suppressed: 790/4447; Not driven: 2422/4447. (D) Mean auROC value of all LiCl-activated neurons in each mouse (n = 8 mice) for saline injection and for LiCl injection. Dashed line at auROC = 0.5 indicates no response. Each dot represents the mean auROC value from one mouse, and values for saline and LiCl are connected for a given mouse. ***P = 6.8 x 10^-7^, paired t test. (E) Left: Heatmap showing the response of all Ensure-activated neurons (n = 1588 neurons from 9 mice) to Ensure consumption, sorted by Ensure response onset time. Right: Heatmap showing the response of the same neurons to LiCl injection, also sorted by Ensure response onset time. Horizontal dashed line indicates the division between Early (green) and Late (purple) Ensure neurons. (F) Mean auROC value for LiCl injection for Early Ensure neurons, Late Ensure neurons, and neurons not driven by Ensure. Each dot is the mean from an individual mouse and values for a given mouse are connected by a line. Early-Late: **P = 0.0094; Early-Not Driven: P = 0.16; Late-Not Driven: P = 0.29, paired t test, Bonferroni-corrected post hoc comparisons. (G) Percent of neurons that are Late Ensure for the LiCl & Ensure activated group (54.6% Late) vs. the group only activated by Ensure (18.4% Late). ***P << 0.0001, two-proportion Z test. Error-bars are +/- standard error of the proportion. (H) Schematic for analysis comparing stimulus responses in awake mice to those in anesthetized mice. (I) Heatmap showing the response of all recorded neurons (n = 3625 from 8 mice) to balloon-mediated stomach stretch in the anesthetized session. Neurons are sorted by the mean in the first 9 seconds post balloon inflation. 0-3 s: inflation from 0 to 0.45 mL; 3-17 s: balloon held at 0.45 mL; 17-20 s: deflation from 0.45 to 0 mL. (J) Left: Heatmap showing the response to stomach stretch of neurons found on both the awake and anesthetized imaging days. Right: Heatmap showing the response of the same neurons to LiCl injection during the awake recording day. Neurons are sorted by the mean response to stomach stretch in both heatmaps (n = 1517 neurons from 7 mice). (K) Mean auROC value for LiCl injection of neurons not driven or driven by stomach stretch. Each dot is the mean from an individual mouse and values for a given mouse are connected by a line (n = 8 mice). ***P = 9.6 x 10^-5^, paired t test. (L) Percent of neurons driven by stomach stretch for the LiCl-activated group (51.9% stomach-driven) vs. for the non LiCl-activated group (15.2% stomach-driven). ***P << 0.0001, two-proportion Z test. Error-bars are +/- standard error of the proportion. (M) Left: Heatmap showing the response of Ensure-activated neurons found on both the awake and anesthetized days to Ensure consumption (n = 450 neurons from 7 mice). Right: Heatmap showing the response of the same neurons to stomach stretch. Neurons are sorted by Ensure onset time in both heatmaps. Horizontal dashed line indicates the division between Early (green) and Late (purple) Ensure neurons. (N) Mean response to stomach stretch of Early Ensure neurons, Late Ensure neurons, and neurons not driven by Ensure. Each dot is the mean from an individual mouse and values for a given mouse are connected by a line. Dashed line at Z=0 indicates no response. No comparisons are significant (P > 1). (O) Percent of neurons that are Late Ensure for the stomach stretch & Ensure activated group (53.6% Late) vs. the group only activated by Ensure (23.2% Late). ***P << 0.0001, two-proportion Z test. Error-bars are +/- standard error of the proportion. For all panels, data is displayed as mean +/-SEM unless otherwise indicated.

The day after the above experiments, we anesthetized mice with isoflurane and implanted a balloon in the stomach to record gastric stretch responses **(Figure 3H**, see Methods). During this imaging session, we repeatedly expanded the balloon to 0.45 mL (representing a mostly full stomach volume^46^) over 3 s, followed by a 14 s hold and subsequent deflation to baseline over 3 s **(Figure 3I).** Many neurons were responsive to stomach stretch, with most neurons showing sustained increases that lasted for the duration of the balloon inflation, and a smaller set exhibiting a transient response to stomach stretch **(Figures 3I and S3C)**.

To compare a single neuron’s response to stomach stretch, LiCl, and Ensure, we identified the same neurons across these two days of recording (**Figure 3H**; see Methods). Neurons activated by stomach stretch had, on average, larger magnitude responses to LiCl injection than those not activated by stomach stretch **(Figures 3J-K)**. Stomach stretch-driven neurons also made up a larger fraction of neurons activated by LiCl than expected by chance **(Figure 3L).** This result aligns with the known LiCl-evoked slowing of gastric motility, which leads to increased stomach stretch and likely activation of stretch-sensitive stomach-innervating vagal afferents^47–49^. Similarly, Late Ensure neurons made up a larger fraction of neurons activated by both stomach stretch and Ensure (53.6% vs. 23.2% for neurons only activated by Ensure, **Figures 3M, O**), though the average stomach stretch response was of similar magnitude for Early Ensure, Late Ensure, and non-Ensure-driven neurons **(Figure 3N).** These results are consistent with the hypothesis that Late Ensure neurons sense post-ingestive information such as visceral malaise and mechanical signals from the stomach.

To further test this hypothesis, we also delivered i.p. injections of cholecystokinin (CCK, 10 ug/kg), an endogenous satiety hormone whose effects are mediated by vagal afferents that sense stomach stretch^48,50^ **(Figure S3D-G).** Late Ensure neurons were significantly overrepresented among neurons driven by both CCK and Ensure **(Figure S3H)**, although there was no difference in mean population response to CCK across Early Ensure, Late Ensure, and non-Ensure-driven neurons **(Figure S3I)**. In a separate set of experiments, we delivered LiCl injections on the first day of recordings, followed by CCK on the second day, and examined neurons recorded across both sessions **(Figure S3J).** As with LiCl, we observed many cells with robust increases or decreases in activity in response to CCK, although with a much faster onset time than for LiCl injections **(Figure S3D-G).** The response to CCK was also sustained for at least 25 minutes post injection. Finally, neurons exhibited mixed sensitivities to LiCl and/or CCK, with 25% of neurons driven by both stimuli, 27% driven only by LiCl, and 13% driven only by CCK **(Figure S3K-L).** Together, these findings demonstrate functional specialization among LPBN neurons responding to various post-ingestive GI stimuli.

### LPBN neurons also track other behavioral variables

In addition to feeding-related stimuli, LPBN responds to various other types of interoceptive signals and modalities including temperature, pain, and general alarm states^3,4^. Thus, we next asked how factors such as somatic pain – and associated changes in locomotion and arousal – affect the recorded LPBN neurons, and how these responses compare to feeding-related activity. To this end, we delivered mild tail shocks (2 x 5 ms pulses over 50 ms, 0.3 mA), which are known to activate LPBN^20,51^. In these sessions, tail shocks were randomly interleaved with Ensure presentations, with a 20 s ITI between stimuli as above **(Figures 3A and 4A)**. We found that many LPBN neurons were activated by shock onsets, with responses lasting from less than 1 s to ∼5 s **(Figure 4Bi)**.

**Figure 4.**
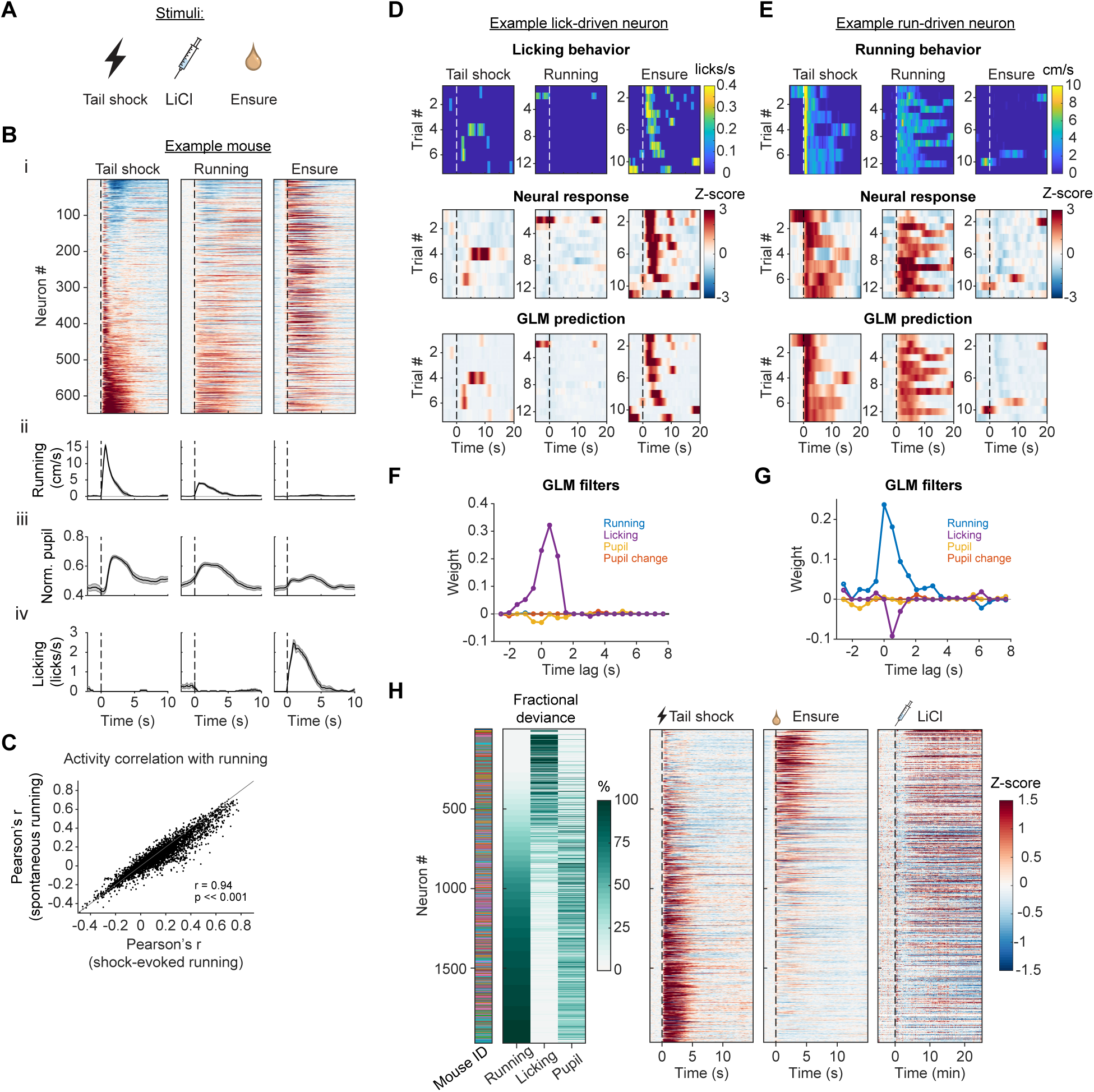
Behavioral variables help explain the activity of LPBN neurons. (A) Stimuli delivered during imaging sessions. (B) i: Heatmaps showing the mean response of LPBN neurons (n = 630) from one example mouse to tail shock (n = 20 trials), running bouts during the ITI (n = 24 bouts), and Ensure consumption (n = 40 trials). ii: Corresponding average traces of running speed during the three stimuli. iii: Average normalized pupil area during the three stimuli. iv: Average lick rate during the three stimuli. Error bars are +/- SEM. (C) Correspondence between activity correlation with running bouts and activity correlation with shock-evoked running across all neurons (Pearson’s correlation; n = 3348 neurons from 9 mice). (D) Top row: Heatmap of licking rate (one row per trial) for an example mouse during tail shock, running bouts, and Ensure delivery. Middle row: Heatmap response of an example neuron whose activity is closely tied to licking. Bottom row: Corresponding GLM prediction of the neuron’s activity based on GLM filter weights for running, licking, pupil, and pupil changes. (E) Same as (D) for a neuron well-driven by running bouts. Top row displays running speed instead of licking. (F) GLM filters showing the influence of each behavioral variable on the activity of the neuron in (D). (G) GLM filters showing the influence of each behavioral variable on the activity of the neuron in (E). (H) Left: Heatmap showing the fractional deviance of running, licking, and pupil area for each neuron (1966 neurons from 9 mice). Right: Heatmap of the average response of these neurons to tail shock, Ensure, and LiCl. In both heatmaps, neurons are sorted by values for running fractional deviance. Consistent with Figure 3, neurons that are well-driven by Ensure and LiCl also show strong fractional deviance for licking.

As expected from previous studies^52^, tail shock stimuli also elicited brief increases in running and pupil dilation (an index of arousal; **Figure 4Bii-iii).** Thus, the shock-evoked changes in activity may be in response to somatic pain, shock-evoked running or arousal, and/or some other bodily changes. To examine these possibilities, we tested how running influences LPBN activity. We searched for running bouts (defined as a minimum speed of 1 cm/s for at least 1 s, preceded by 1 s of immobility) during periods not associated with any stimulus (either during baseline recordings or from 6-19 s after stimulus offset within the ITI). We then computed average run-bout-triggered responses. Surprisingly, we observed a strong correspondence between the neurons activated during spontaneous running bouts and those driven by tail shock **(Figures 4B-C and S4A)**. This finding indicates that at least part of the shock-evoked activity may be due to the running induced by tail shock rather than the shock itself. In fact, including running bouts as a potential driver of neural activity during the session (in addition to two other variables, Ensure and LiCl) increased the percentage of imaged neurons activated by at least one variable from 60% to 91% (1190/1966 to 1800/1966). This strong correlation between activity of a subset of LPBN neurons and running demonstrates the importance of tracking behavioral states during LPBN recordings.

To more robustly and accurately dissociate the effects of running and other behavioral variables that modulate LPBN neuron activity, we implemented a generalized linear model (GLM) that quantifies how much each behavioral variable can explain the activity of individual neurons^53,54^. We included four predictors in the model: running speed, lick rate, pupil area, and change in pupil area (i.e., constriction/dilation, which provides complementary information about arousal states^43^). We also allowed for multiple delays between each predictor and its effect on neural activity (see Methods). The model captured the diversity of responses to presented stimuli, for example by revealing an early or delayed influence of licking for Early and Late Ensure neurons, respectively **(Figure S4C).** Several different types of response profiles also emerged from this analysis, including neurons influenced solely by running or licking **(Figure 4D-G).** For instance, **Figure 4D** shows an example lick-driven neuron, with variable activity across trials that closely tracks the timing and rate of licking. Note that a strong GLM filter for licking (e.g., **Figure 4F)** does not indicate that the neural activity is directly caused by licking behavior. Licking a drop of Ensure could be associated with several events or stimuli whose timing is locked to licking, from motor preparation before licking, to taste and other oral sensations during licking, to food arriving at parts of the GI tract with stereotyped delays after licking **(Figure 1)**; a model with large weights for licking at various delays can reflect any of these associations.

We next used different model iterations to test which behavioral variables are sufficient to explain activity of each recorded neuron in LPBN. To this end, we re-ran the GLM using only one behavioral variable at a time and report the performance of each model as a fraction of the deviance explained under the full model with all variables^54^; average deviance explained for full model across neurons: 0.26 +/- 0.14, mean +/- SD; **Figures S4B and 4H).** This analysis confirmed that many recorded neurons are primarily driven by running (60%, 1123/1883) and form a mostly separate group from neurons strongly driven by licking or Ensure consumption (26%, 494/1883). In contrast, only a minority of neurons showed activity best explained by changes in pupil-indexed arousal (pupil: 10%, 180/1883; change in pupil: 5%, 86/1883). Interestingly, this analysis revealed that Early Transient Ensure neurons **(Figure 2F, H)** did not exhibit activity explained by licking, but were instead sensitive to locomotion **(Figure S4D),** indicating that these neurons’ Ensure responses are likely to due to postural adjustments as the mouse begins to consume Ensure. The Early Sustained Ensure and Late Ensure groups showed mixed influences of licking and running **(Figure S4D)**, suggesting that many neurons may represent licking state, stimuli in the mouth and upper GI tract, and/or gastric and post-ingestive signals, while other neurons may respond to motor behaviors during a trial. These results demonstrate that a diverse repertoire of stimuli and internal states can influence activity in different LPBN neurons in response to the same action – licking for Ensure. Overall, these analyses reveal two somewhat separate functional classes in LPBN: one driven by locomotion and one driven by licking.

The above findings highlight the diversity of functional properties in LPBN across neurons. As a first step towards linking these functional properties to genetically defined cell types in LPBN, we ran the identical experimental protocols from **Figures 2-4** during GCaMP fiber photometry recordings from a particularly well-studied LPBN cell type, CGRP neurons. We chose CGRP neurons because they are concentrated in PBNel (which is encompassed by our FOVs, **Figure 2C)**, and are known to be important in regulating feeding, visceral malaise, pain, and arousal^3,6,11,20,55^. We expressed Cre-dependent GCaMP (DIO-GCaMP6s) in the LPBN of *Calca-Cre* mice and recorded bulk fluorescence through an optic fiber implanted above LPBN **(Figure S8A).** As in **Figure 3** and previous Fos studies^6,45^, CGRP neurons responded to LiCl injection with a strong activation for tens of minutes post injection **(Figure S8B)**. We next presented drops of Ensure and tail shocks as in our previous experiments while tracking running speed, lick rate, and pupil area. CGRP neurons were robustly activated by consumption of drops of Ensure, and showed variable activity driven by tail shock and running bouts **(Figure S8C-E)**. Applying the same GLM analysis to these signals revealed that bulk CGRP neuron activity is best explained by licking and Ensure consumption rather than running **(Figure S8F-H)**, consistent with our findings that LiCl-driven neurons also are more commonly driven by licking (with sustained duration and/or late onset) than by running **(Figure 4H).** In summary, we observe an emerging functional specialization of neurons in LPBN, many of which respond to behavioral variables – in particular running – while others respond more to licking and/or interoceptive stimuli.

### Spatial organization of functional subtypes in LPBN

Our findings reveal groups of LPBN neurons that are sensitive to different stimuli (for example, activation following LiCl injections or during locomotion) and that show distinct response kinetics on the timescale of seconds (for example, Early vs. Late Ensure). We next asked whether these groups are spatially organized within LPBN. There are several lines of evidence suggesting a functional topography in this region. First, genetic subtypes are spatially clustered within LPBN subregions on the scale of several hundred micrometers^22,23^. Second, the NTS, a region upstream of LPBN, exhibits coarse viscerotopy in anesthetized, in vivo recordings^28^. This viscerotopic map in NTS may be passed on to the LPBN via topographically organized inputs^56^. Finally, several studies have observed spatial clustering of Fos expression in LPBN after different kinds of stimuli^18,35,57–59^. However, examining the viscerotopic organization in LPBN has proven difficult because of the challenges with imaging activity in LPBN *in vivo* with a large enough FOV. We took advantage of our 1 mm GRIN lens preparation to determine how neurons with different functional properties might be spatially organized.

The first indication of such functional clustering came from observing the response of neurons across the FOV to the consumption of Ensure. In the example mice in **Video S5** and **Figure 5A** (top row), Early Ensure neurons tended to be located more posterior and lateral than Late Ensure neurons. To further quantify this spatial clustering, we estimated the probability that two Ensure-driven neurons within a certain distance of each other were both Late Ensure neurons **(Figure 5B).** 6/9 mice showed significant clustering of Late Ensure neurons separate from Early Ensure neurons at a spatial scale of <100 µm as compared to shuffle controls **(Figure S5A).** Consistent with the fact that Late Ensure neurons were more commonly driven by LiCl and stomach stretch **(Figure 3E, G, M, O),** we found that neurons activated by LiCl (including both Ensure-activated and non-activated subsets) are concentrated in the anterior FOV regions near the Late Ensure neurons (**Figure 5A-B**, second row; 9/9 mice showed clustering for LiCl activation above chance, **Figure S5B**). Neurons activated by stomach stretch also exhibited spatial clustering at the 100 µm scale in 7/7 mice **(Figure S5D-F).** Similarly, neurons driven by running showed significant clustering (8/9 mice; **Figure 5A-B** bottom row and **Figure S5C),** typically in more posterior regions, away from cells responsive to licking or LiCl.

**Figure 5.**
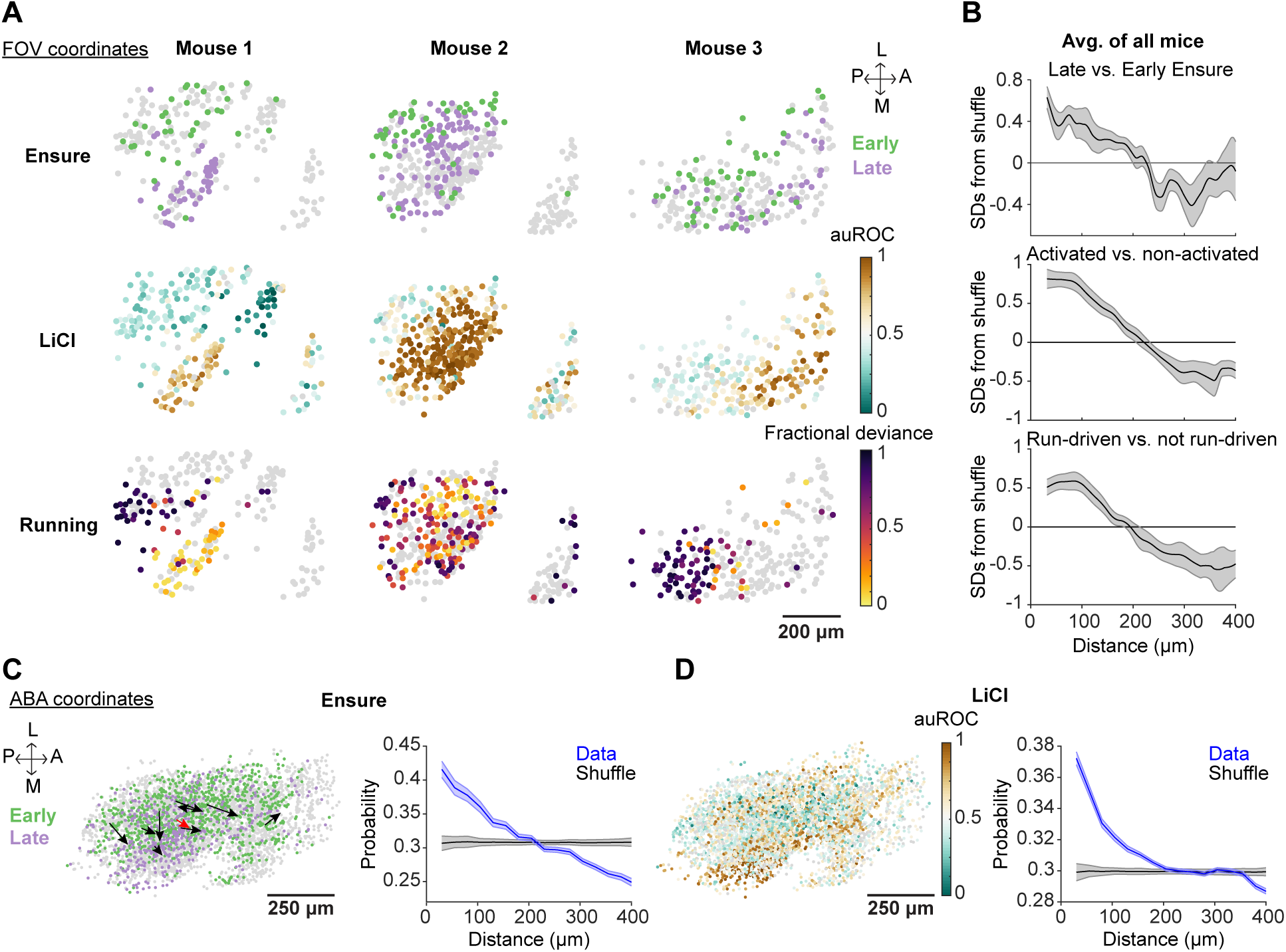
Spatial organization of LPBN functional groups. (A) Cell masks in field of view (FOV) coordinates for three example mice, colored by Early or Late Ensure (top row), LiCl response (middle row), and modulation by running (bottom row). (B) Likelihood that a pair of neurons spaced different distances apart were both Late Ensure neurons (expressed as the number of standard deviations [SDs] from shuffle controls), averaged over all mice (n = 9; top row). Middle and bottom rows show the spatial clustering for LiCl-activated neurons (vs. non-activated neurons) and neurons driven by running (vs. not driven by running), respectively. Shaded regions show SEM across mice. See **Figure S5** for statistical analyses. (C) Left: Cell masks (n = 3602 from 9 mice) in Allen Brain Atlas coordinates, colored by Early Ensure or Late Ensure. Neurons not driven during consumption are labeled in gray. Arrows indicate the average direction from Early to Late neurons for each mouse (red: average across all mice). Right: Likelihood that a pair of neurons spaced different distances apart were both Late Ensure. Shaded regions show SEM across neurons. (D) Left: Cell masks (n = 3602 from 9 mice) in Allen Brain Atlas coordinates colored by LiCl response. Right: Likelihood that a pair of neurons spaced different distances apart were both activated by LiCl. Shaded regions show SEM across neurons.

Small variations in GRIN lens placement may result in sampling slightly different subregions of LPBN across mice. To determine whether imaging location underlies variability in functional maps across mice, we used post hoc coronal sections containing the GRIN lens tract to align our FOVs to the Allen Mouse Brain Common Coordinate Framework and plot all neuron locations in this coordinate system **(Figures 2C and S2A-B).** While this method of alignment is somewhat coarse, we observed consistent global organization across mice for the stimuli presented **(Figures 5C-D and S5G-H).** For instance, in this coordinate system, the vector between the average location of Early Ensure neurons to that of Late Ensure neurons progresses in the same posterior to anterior direction (**Figure 5C**; 8/9 mice) and, to a lesser extent, from lateral to medial (7/9 mice). In addition, we observed a hot spot of LiCl-active neurons in a subregion that putatively aligns with PBNel CGRP neurons **(Figure 5D),** consistent with previous work^6,35,44,45^ and our photometry experiments showing CGRP activation in response to LiCl **(Figure S8B).** Overall, these results demonstrate a spatial clustering of functional subtypes in LPBN, which could map onto genetic cell types and/or onto different anatomical inputs to LPBN.

### Slow changes in LPBN activity over tens of minutes during a meal

So far, we have only considered LPBN activity on the fast, seconds-long timescale. However, we observed in **Figure 1H-I** that during a meal, the stomach size slowly increases over the course of many minutes. Furthermore, brain regions upstream (NTS) and downstream (insular cortex, InsCtx) of LPBN have been shown to encode satiety through slow, minutes-long changes in ongoing activity over the course of a meal^60–62^. Thus, we next asked whether LPBN neurons also show minutes-long changes in ongoing activity during feeding sessions.

To answer this question, we first concatenated activity from the last 10 s of each 20-s inter-trial interval (ITI) following Ensure delivery, similar to our earlier studies of InsCtx **(Figure 6A)**^60^. We used this period to avoid contamination from any stimulus-induced activity from the previous trial, and we excluded periods involving licking or running bouts. Following forty equally spaced trials of Ensure delivery (20 s per trial), we also recorded a 30-minute free-feeding protocol in which mice had continuous access to Ensure, and we included baseline periods of quiet waking before and after this free-feeding protocol **(Figure 6A)**. Mice ceased licking towards the end of the free-feeding protocol, indicating subjective satiation. We observed many neurons that showed either a gradual increase or decrease in ongoing activity over the course of this hour-long imaging session **(Figures 6B-C).** Notably, the degree and sign of slow activity changes in individual neurons (quantified by mean GCaMP8s fluorescence in the last two minutes of quiet waking activity in the session minus the first two minutes) were very similar over two consecutive imaging sessions using the identical protocols (1505/2897 neurons, 52%, had a significant change in the same direction on both days; **Figures 6C-D)**. This consistency across days suggests that the slow changes likely were due to real variations in neural activity as opposed to technical artifacts, such as a drift in the imaging FOV over time.

**Figure 6.**
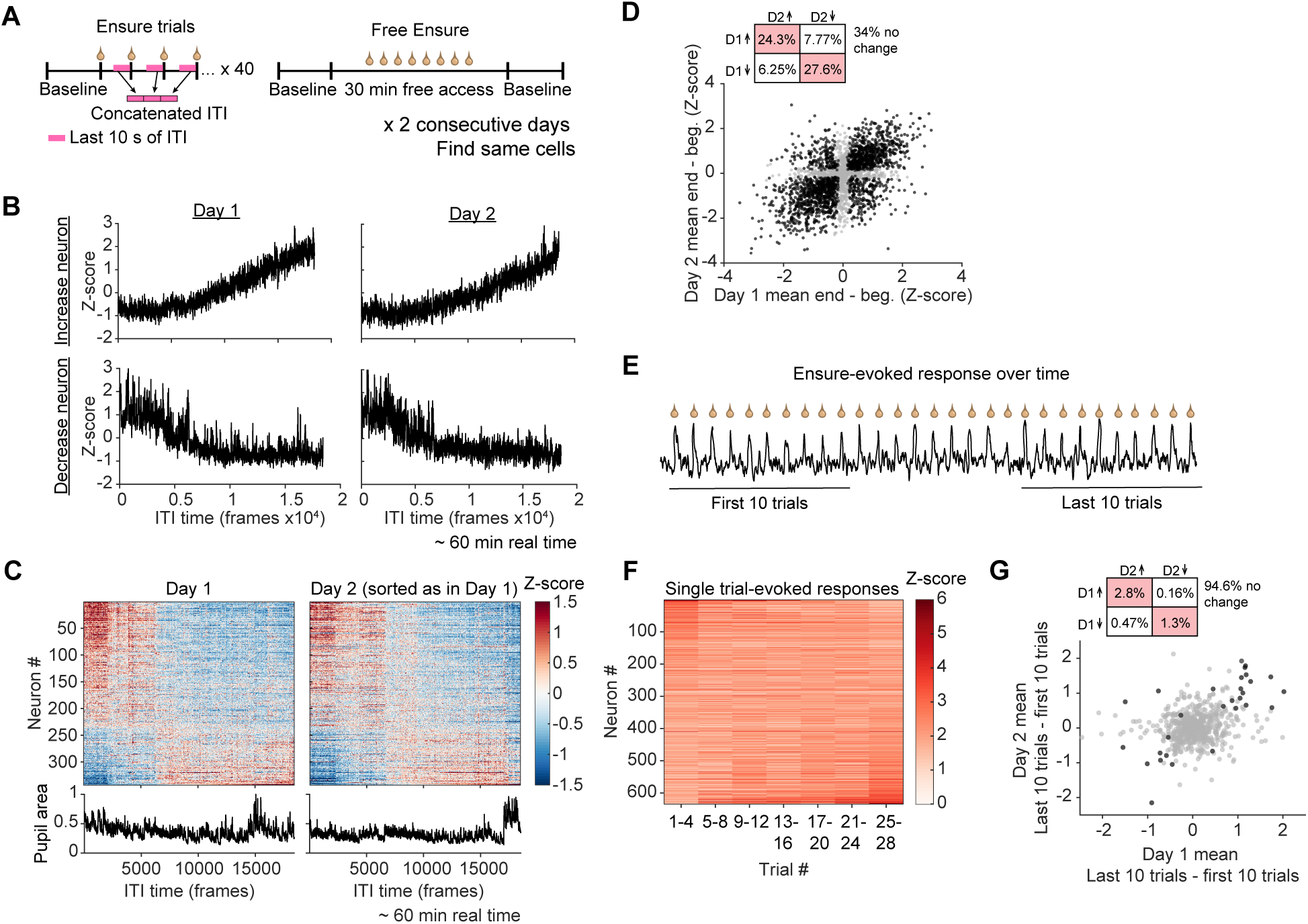
Slow changes in LPBN activity over tens of minutes. (A) Schematic of experiment for analyzing slow activity changes during baseline periods and late-ITI periods (pink) without licking or locomotion. (B) Slow activity changes on two consecutive days for an example neuron that increases its activity (top) and an example neuron that decreases its activity (bottom) throughout the recording session. (C) Top: Heatmaps showing the concatenated ITI and quiet-waking activity of single neurons on two consecutive days of imaging from an example mouse. Neurons are sorted by mean activity in the last 2 minutes – mean activity in the first 2 minutes of the session on Day 1. Bottom: Concatenated pupil area (normalized to maximum within the day) from Days 1 and 2. (D) Scatter of the change in activity (mean of last 2 min – mean of first 2 min) on Day 1 with that on Day 2 across all recorded neurons found on both days (n = 2897 neurons from 8 mice). Black dots indicate neurons that had a significant change in activity on both days (P < 0.05, see Methods). Gray dots indicate neurons that did not have a significant change on at least one day. Inset: Percent of neurons found on both days that had a significant increase (τ) or decrease (,) in activity. D1: day 1; D2: day 2. (E) Example activity trace of a neuron responding to single Ensure trials (20 s ITI). (F) Heatmap showing the maximum Ensure response, averaged for every 4 consecutive trials, of neurons significantly activated by Ensure on 2 consecutive days (n = 634 neurons from 8 mice). Neurons are sorted by the difference between the values of the last column and first column. (G) Scatter between Day 1 and Day 2 values of the difference between the mean of the maximum Ensure responses across the last 10 trials and the mean of the maximum Ensure responses across the first 10 trials (for Ensure-activated neurons found on both days; n = 634 neurons from 8 mice). Black dots are neurons that had a significant difference in this metric on both days (P < 0.05, see Methods). Gray dots are neurons that were not significant on at least one day. Inset: Percent of Ensure-activated neurons found on both days that had a significant increase (τ) or decrease (,) in Ensure response. D1: day 1; D2: day 2.

Next, we asked whether a neuron’s slow change in activity was related to the dynamics of its Ensure-evoked response on the faster timescale of seconds. For example, we might expect Late Ensure neurons, which we hypothesize to be responsive to stomach stretch **(Figure 3M, O)**, to increase their tonic activity as stomach volume increases over the session **(Figure 1H-I).** To explore this question, first we sorted the slow change in activity of all Ensure-activated neurons by their average Ensure response onset times **(Figure S6A).** We observed no consistent relationship between the temporal dynamics of responses to a drop of Ensure (i.e., Early vs. Late onset) and slow changes in activity **(Figure S6A-B)**. This suggests that the patterns of LPBN neurons that change their activity at fast (seconds) and slow (minutes-hours) timescales during feeding are distinct from one another. Though we cannot be certain that these slow changes were due to feeding alone, we observed no slow change in pupil-indexed arousal over the course of the sessions **(Figure S6C)**.

Satiety might also drive across-meal changes in the magnitude of a neuron’s acute response to feeding, as has been observed in the hypothalamus **(Figure 6E)**^63^. To determine how an Ensure-activated neuron’s response to consuming a drop of Ensure changes as a mouse becomes sated, we averaged the peak response of every four trials throughout the session. While some neurons showed gradual or abrupt increases or decreases in peak response over 28 trials, other neuron responses were quite stable **(Figure 6F)**. We then averaged each neuron’s peak response across the first 10 Ensure trials of the day and compared it to the average peak from the last 10 Ensure trials of the day. We found that only 4.1% (26/634) of Ensure-driven neurons showed a significant, within-session change in feeding-evoked response magnitude that was consistent across two days of imaging **(Figures 6G, S6D).** This finding suggests that most LPBN neurons encode consumption bouts in a largely stable manner over the course of a meal, while a small minority of LPBN neurons may encode satiation by changing their response to consumption bouts throughout a meal.

### Insular cortex input to LPBN activates post-ingestive LPBN neurons

As a sensory hub, LPBN receives input from several brain regions, including bottom-up input from the periphery (spinal cord) and other brainstem areas (NTS, AP, trigeminal nuclei) and top-down input from forebrain regions, such as the hypothalamus, amygdala, and cortex^2,64,65^. In some cases, these inputs have been shown to target specific subregions and genetically defined neural populations within LPBN^16,56,65–68^. Thus, we hypothesized that different input streams may differentially target our functionally defined LPBN neuron populations. We tested this possibility by focusing on the dense, excitatory projection from InsCtx neurons to LPBN^65,66^. Posterior granular InsCtx is the primary viscerosensory cortex and contains a viscerotopic map of internal organs (e.g., GI tract, heart, lungs)^69–71^. Further, InsCtx activity appears to encode current as well as future physiological states (e.g., hunger/satiety)^60,72^, suggesting the potential for anticipatory regulation of LPBN by InsCtx. To obtain initial clues as to how cortical input regulates LPBN, we examined whether InsCtx inputs to LPBN (InsCtx^LPBN^) were sufficient to drive LPBN activity and whether this drive was restricted to specific subsets of functionally or spatially defined neurons.

To study the influence of InsCtx^LPBN^ inputs on LPBN neuron activity, we expressed AAV1-ChrimsonR-tdTomato throughout InsCtx and prepared LPBN for two-photon calcium imaging as described in **Figure 2 (Figures 7A-C and S7A)**. To optogenetically stimulate InsCtx^LPBN^ axons, we delivered red light (620 nm, 2.5 mW, 8 ms pulses at 15.6 Hz for 2 s) through the GRIN lens while simultaneously recording ribo-GCaMP8s signals from LPBN neurons. We found that InsCtx^LPBN^ axon stimulation potently activated LPBN neurons in a consistent manner across trials and did not induce any obvious behavioral effects **(Figures 7D-E and S7B-C)**. This finding suggests that the InsCtx input alone can drive robust activity in LPBN. In addition, despite extensive ChrimsonR expression throughout InsCtx **(Figure S7A)** and broad innervation of LPBN by ChrimsonR-tdTomato-expressing InsCtx^LPBN^ axons **(Figure 7B),** InsCtx^LPBN^ axon stimulation activated only a subset of recorded LPBN neurons (29.8%, 481/1614).

**Figure 7.**
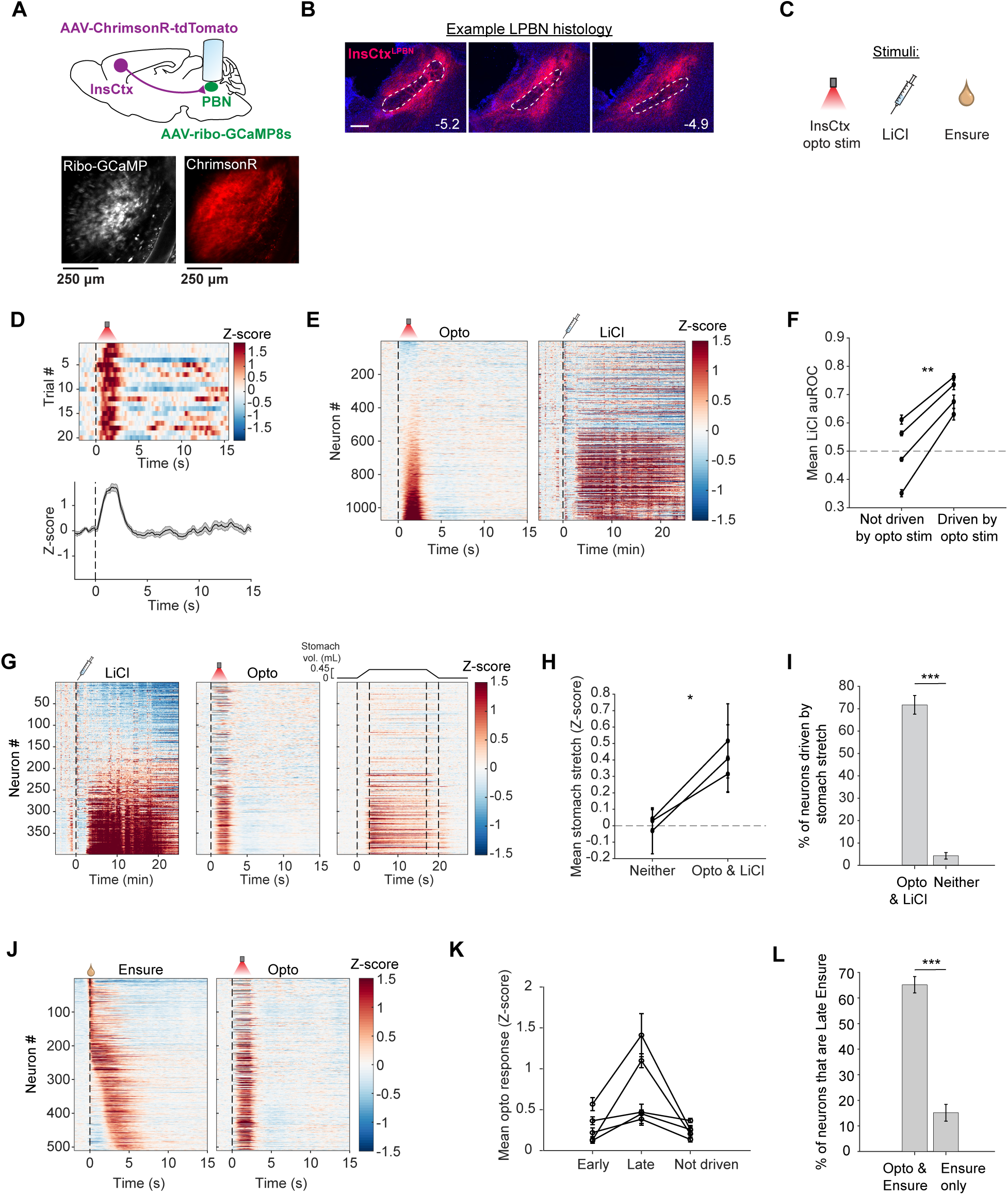
InsCtx^LPBN^ axons activate post-ingestive, LiCl-driven LPBN neurons. (A) Optogenetic stimulation of InsCtx^LPBN^ axons during two-photon calcium imaging of LPBN. Top: Schematic for ChrimsonR expression in InsCtx and ribo-GCaMP8s expression in LPBN with GRIN lens implantation above LPBN. Bottom: Example two-photon FOV showing ribo-GCaMP8s expression in LPBN neurons (left) and ChrimsonR expression in InsCtx^LPBN^ axons (right). (B) Coronal histology showing expression pattern of InsCtx^LPBN^ axons throughout LPBN in an example mouse. Dashed white lines outline the superior cerebellar peduncle. Scale bar: 250 μm. (C) Stimuli delivered during imaging sessions. (D) Example neuron driven by InsCtx^LPBN^ stimulation. Top: Heatmap showing the neuron’s response to single 2 s stimulation trials. Bottom: Mean activity trace (n = 20 trials) aligned to stimulation onset at 0 s. (E) Left: Heatmap showing the mean response (n = 20 trials) of single LPBN neurons (n = 1074 neurons from 4 mice) to optogenetic photostimulation of InsCtx^LPBN^ axons (“opto”). Stimulation occurs from 0-2 s. Right: Heatmap showing the response of the same neurons to LiCl injection. For both heatmaps, neurons are sorted by the mean Z-score from 0-2 s during photostimulation. (F) Mean auROC value for LiCl injection of neurons not driven (left) or driven (right) by InsCtx^LPBN^ stimulation. Each dot is the mean from an individual mouse and values for a given mouse are connected by a line. **P = 0.0055, paired t test. (G) Left: Heatmap showing the response of neurons found on both the awake and anesthetized imaging days to LiCl injection (n = 399 neurons from 3 mice). Middle: Heatmap showing the response of the same neurons to InsCtx^LPBN^ photostimulation. Right: Heatmap showing the response of the same neurons to stomach stretch. For all three heatmaps, neurons are sorted by auROC estimates of LiCl-evoked responses. (H) Mean response to stomach stretch of neurons that were not driven by InsCtx^LPBN^ stimulation or LiCl vs. neurons driven by both InsCtx^LPBN^ stimulation and LiCl. Each dot is the mean from an individual mouse and values for a given mouse are connected by a line. Dashed line at Z = 0 indicates no response. *P = 0.0207, paired t test. (I) Percent of neurons recorded on both days that are driven by stomach stretch for the InsCtx^LPBN^ opto & LiCl-activated group (71.6%) and for the group not driven by opto or LiCl (4.1%). ***P << 0.0001, two-proportion Z test. Error-bars are +/- standard error of the proportion. (J) Left: Heatmap showing the response of all recorded Ensure-activated neurons to Ensure consumption in experiments involving InsCtx^LPBN^ opto (n = 510 neurons from 5 mice). Right: Heatmap showing the response of the same neurons to InsCtx^LPBN^ stimulation. For both heatmaps, neurons are sorted by Ensure response onset time. (K) Mean response to InsCtx^LPBN^ photostimulation of Early Ensure neurons, Late Ensure neurons, and neurons not driven by Ensure. Each dot is the mean from an individual mouse and values for a given mouse are connected by a line. Early-Late: P = 0.16; Early-Not Driven: P > 1; Late-Not Driven: P = 0.21, paired t test, Bonferroni-corrected post hoc comparisons. (L) Percent of Ensure-responsive neurons classified as Late onset for the InsCtx^LPBN^ opto & Ensure-activated group (65.2%) and for the group only activated by Ensure (15.1%). ***P << 0.0001, two-proportion Z test. Error-bars are +/- standard error of the proportion. For all panels, data is displayed as mean +/-SEM unless otherwise indicated.

InsCtx-driven LPBN neurons had, on average, larger responses to LiCl injection than those not driven by InsCtx **(Figures 7E-F and S7C)**. In a subset of mice (n=3) where we also examined responses to stomach stretch, we observed that neurons driven by both InsCtx^LPBN^ and LiCl also had larger mean responses to stomach stretch than those driven by neither of these two stimuli **(Figures 7G-H and S7D-E).** Further, a larger fraction of the subgroup driven by both InsCtx^LPBN^ and LiCl was also driven by stomach stretch compared to neurons not driven by LiCl or InsCtx^LPBN^ (71.6% vs. 4.1%; **Figure 7I)**. Consistent with our previous finding that Late Ensure neurons were overrepresented in the LiCl or stomach stretch-driven groups **(Figure 3G, O)**, Late Ensure neurons also made up a larger proportion of the LPBN subgroup driven by InsCtx (68.7% vs. 48.1% of the Ensure only group; **Figures 7J, L and S7F**). Finally, in some mice but not others, InsCtx^LPBN^ axon stimulation more strongly drove Late Ensure neurons than Early Ensure neurons **(Figures 7K and S7F)**. Together, these data indicate that InsCtx specifically targets LPBN neurons that encode stomach stretch and/or other post-oral GI stimuli, hinting at a potential role for InsCtx input to LPBN in regulating appetite suppression.

Consistent with the above findings and with previous work, we observed a particularly high density of InsCtx^LPBN^ axons in the PBNel region, where CGRP neurons are found **(Figures 7B and S7G)**^66,68^. Given that both LiCl^6,45^ and stomach stretch can activate CGRP neurons **(Figure S8B, I-J)**, we next asked whether CGRP neurons are driven by InsCtx^LPBN^ axon stimulation. To answer this question, we expressed ChrimsonR-tdTomato throughout InsCtx of *Calca-Cre* mice and performed fiber photometry recordings from GCaMP6s-expressing CGRP neurons in LPBN as previously described **(Figure S7G).** In response to 2 s of InsCtx^LPBN^ axon photostimulation, we observed a robust increase in CGRP GCaMP fluorescence in 5/5 mice, indicating that InsCtx can drive activity in CGRP neurons **(Figures S7H-I)**. Thus, we predict that CGRP neurons are a subset of the LPBN subpopulation driven by InsCtx^LPBN^ input, LiCl, and stomach stretch, raising the possibility that InsCtx feedback to LPBN may support anticipation of future visceral changes.

## Discussion

LPBN is a key integrator of multimodal body signals and regulates physiological and behavioral responses related to appetite, fluid balance, breathing, and pain. These functions make characterizing its sensory coding an important step to understanding many interoceptive sensory processes. However, unique challenges have hampered the study of LPBN and other circuits underlying body-brain communication during natural behaviors such as feeding. First, we lacked a detailed understanding of the real-time dynamics of food movement through the GI tract. Second, we lacked strategies to tease apart various contributions to an LPBN neuron’s activity, given the partially correlated changes in interoceptive signals (e.g., food movement through the GI tract) and exteroceptive signals (e.g., body movement) during feeding behavior. Finally, some physiological effects are not reversible for hours, preventing repeated stimulation and extensive characterization of functional properties of neurons.

To begin to overcome these challenges, we first monitored the movement of food through the upper GI tract during feeding by adapting X-ray videofluoroscopy for awake, head-fixed mice. In parallel experiments, we combined large-scale recordings of LPBN across days with visceral and feeding-related stimuli, behavioral state tracking, and statistical modeling to reveal interoceptive and other sensitivities of LPBN neurons. Below, we discuss our findings of functionally specialized and spatially clustered groups of neurons that encode ingestion events and other body information, including oral or post-ingestive feeding-related signals as well as locomotion and arousal.

### GI signals contributing to the activity of Early and Late Ensure neurons

We found that LPBN neurons responded to food consumption with varied temporal dynamics, suggesting that they encode distinct processes. Early Ensure neurons became active prior to or at the onset of licking and had a transient or sustained duration, while Late Ensure neurons became active 1-4 s after the onset of licking **(Figures 2E-H)**. Given the similarity of the onset of feeding responses in Late Ensure neurons to the arrival times of food in the esophagus and stomach (but not to arrival in the duodenum, **Figures 1D-E, G, S1A)**, we hypothesize that Late Ensure neurons encode food-related signaling that occurs during or after swallowing, while Early Ensure neurons largely respond to stimuli in the mouth. For example, some Early Ensure neurons may encode taste, as several electrophysiology and Fos studies in rats have demonstrated taste responses throughout LPBN^58,73,74^. Additionally, Satb2-expressing neurons in the ventral-lateral region of the PBN have been shown to respond to multiple taste modalities with a fast onset^26,27^. Some LPBN neurons also respond to tactile stimulation of the tongue^24^ and exhibit activity that is correlated with tongue EMG signals^73^, suggesting that Early Ensure neurons could be sensitive to oral mechanosensory signals. Though our study shows that licking drives many Early Ensure neurons **(Figures 4D, F, H, S4D)**, we cannot distinguish whether this activity is due to taste-associated signals, mechanical signals, or both. We did observe some examples of neurons that became active during ITI licking (when no Ensure was present)^39^; however, most mice did not lick during the ITI. Future work could tease apart this early consumption-related activity by combining recordings during feeding with subsequent delivery of “pure” mechanical or taste stimuli under anesthesia (similar to our experiments involving stomach stretch).

Notably, many Early Ensure neurons became active within the 1 s *preceding* the first lick **(Figure 2G)**, though they reached maximum activation during licking. This pre-lick activity could be due to body or orofacial movements as the mouse prepares to lick (e.g., in some cases, mice stopped running immediately prior to licking). Consistent with this possibility, the activity of most Early Transient neurons was best predicted by locomotion or body movement **(Figure S4D)**. In other cases, this activity may be anticipatory and predictive of the incoming orosensory signals discussed above (e.g., related to the smell of Ensure or the sound of the syringe pump during delivery). Such anticipatory activity has been observed in other feeding-related neurons, such as hypothalamic AgRP neurons, which are inhibited by distal sensory signals associated with food (e.g., sight or smell) prior to the onset of consumption^75–77^.

In contrast, Late Ensure neurons likely encode post-ingestive signals such as gastric stretch. Supporting this hypothesis, we found that neurons driven by stomach stretch, LiCl, or CCK were more likely to be Late Ensure neurons **(Figures 3G, O and S3H)**. LiCl can induce stomach stretch by slowing gastric emptying, particularly when delivered after a meal^47^. Further, GLP1R+ vagal afferents, which are activated by stomach stretch and CCK^48^, partially mediate the LPBN response to LiCl^49^. Given that LiCl, stomach stretch, and CCK signals share a common sensory pathway to LPBN, we propose that this pathway contributes to the activity of Late Ensure neurons during food consumption. Indeed, a previous study showed that the vagus nerve mediates gastric stretch responses of Pdyn-expressing LPBN neurons, though whether these neurons exhibit a late-onset consumption response is not clear^24^. In addition to the vagal pathway, malaise signals such as those caused by LiCl reach LPBN via the area postrema (AP)^67,78,79^. We observed that many neurons not driven by feeding were strongly activated by LiCl **(Figures 3F**, **4H, and 5A)**, suggesting that these neurons may be dedicated to signaling malaise through AP projections to LPBN. In some cases, such neurons were found even more anterior in LPBN than Late Ensure neurons **(Figure 5A)**, consistent with a high density of LiCl-sensitive, GLP1R+ AP axons in anterior regions of LPBN^67^.

Late Ensure neurons exhibited a wide range of onset times from 0.9-4 s after the first lick **(Figure 2G)**, suggesting that they form a heterogenous group. It is possible that neurons with earlier onsets within this range are sensitive to both esophagus and stomach (as has been shown for the Pdyn population^24^), while those with later onsets respond only to entry into the stomach. We did not observe neurons with response onset times aligned to the arrival of food in the duodenum (∼10 s). This may be due to several factors including variable likelihood and timing of arrival across trials and/or earlier, anticipatory responses to upcoming duodenum stimulation. Future combination of calcium imaging or other single-neuron recording methods with fluoroscopy would be challenging but would allow for moment-to-moment correlations between GI transit and neural responses.

### Slow changes in ongoing LPBN neural activity may encode satiety

We observed many LPBN neurons that showed either an increase or a decrease in ongoing (inter-trial) activity across tens of minutes as mice consumed Ensure during a one-hour imaging session, independent of changes in arousal state **(Figures 6B-D and S6C)**. Neurons with these slow activity changes may track cumulative food intake through nutrient absorption and/or changes in stomach volume. Indeed, our X-ray fluoroscopy experiments showed a gradual increase in stomach volume **(Figures 1H-I and S1D)** that occurred on the same minutes-long timescale as the slow changes in neural activity. However, there was no relationship between a neuron’s slow change in activity and the dynamics of its fast-timescale response to Ensure consumption **(Figure S6A-B)**.

These ongoing activity changes in LPBN also echo our previous work showing that InsCtx ongoing activity patterns reflect physiological state, with slow increases or decreases in neural activity as a mouse transitions from hungry to sated^60^. InsCtx receives both direct and indirect (via visceral thalamus) input from LPBN, suggesting that LPBN ongoing activity may also encode physiological state and supply this information to InsCtx^64,66,80^. It is important to note, however, that we only tracked satiety and arousal state; slow changes in activity of certain LPBN neurons may also track other body states, including fluid balance^10,81,82^, which also changes as mice consume liquid food.

In contrast to the changes in ongoing activity throughout a session, we found that Ensure-driven responses of most LPBN neurons were stable across Ensure-consumption trials **(Figures 6F-G)**. LPBN-projecting Prlh/TH NTS neurons similarly do not show state-dependent changes in consumption-evoked activity^61,62,83^. Yet, neurons in other feeding-related areas, such as the paraventricular hypothalamus MC4R neurons (PVH^MC4R^ neurons, some of which project to LPBN to promote satiety^84^), show increased consumption-evoked activity as a mouse becomes sated^63^. Why might responses in some LPBN and NTS neurons remain stable over trials? We speculate that these sensory-related responses provide a steady signal to downstream brain regions to indicate ongoing consumption, while separate systems track satiation.

### Importance of tracking behavioral variables in interoception research

A key component to studying neural activity in awake animals is that sensory stimuli can induce associated body and behavioral state changes. Tracking these secondary effects is particularly important in studies of interoception to dissociate which physiological or brain state changes actually modulate a given neuron’s activity. For example, we show that administering mild tail shocks to head-fixed mice activates certain LPBN neurons while driving concurrent increases in locomotion and pupil-indexed arousal. We found that the activity of many shock-responsive LPBN neurons was highly correlated with and could be well explained by associated changes in locomotion, as spontaneous locomotion drove similar responses **(Figure 4).** Had we not recorded locomotion here, we may have concluded that shock-responsive LPBN neurons encode the painful or surprising effects of tail-shock^20,21^. Instead, we propose that running-linked LPBN responses may stem from spinoparabrachial tract inputs, which carry innocuous tactile signals (along with pain and temperature information)^4,15–17,85^ to LPBN (e.g., footfalls on the running wheel during locomotion). A minority (10%) of LPBN neurons exhibited responses most linked to pupil-indexed arousal, which is smaller than might be expected from previous literature documenting arousal signaling in LPBN^11,13,29^. However, pupil-indexed arousal still correlated strongly with LPBN activity, albeit to a lesser extent than running (e.g., running could explain at least half of the activity in 58% of neurons, while pupil accounted for half of the activity in only 26% of neurons; **Figure 4H)**.

Similarly, we observed some Ensure-responsive neurons whose activity could be best explained by movement **(Figure S4D)**. We hypothesize that these neurons respond to tactile stimuli that result from a mouse adjusting its body posture to ingest Ensure on the wheel. Interestingly, while the activity of most neurons could be statistically accounted for by either licking or running alone, a subset displayed mixed coding **(Figures 4H and S4D)**. These neurons may reflect the integration of signals from multiple pathways in LPBN (e.g., tactile stimuli from spinoparabrachial tract and GI stimuli from the NTS and vagus nerve) to provide a multimodal representation of an animal’s current state.

### LPBN neuron subtypes are spatially organized

We found that neurons with similar functional properties were spatially clustered within LPBN **(Figure 5)**. How might this organization come about? One possibility is that it is inherited from the NTS, where there is an established anatomical and functional organization of inputs from different body organs^28,35,86^. In fact, neurons encoding oral or stomach stimuli are spatially segregated in NTS^28^, similar to our findings for Early vs. Late Ensure neurons in LPBN **(Figure 5A-C)**. The viscerotopy in NTS could be preserved in LPBN through projections from distinct regions of NTS targeting distinct regions of LPBN, as has been shown in rats^56^. LPBN also receives input from other ascending sensory pathways including from the AP, trigeminal ganglia (both directly and via the trigeminal nuclei), and the spinal dorsal horn^4,16,17,78,79,87–89^. Axons from these pathways show distinct (though sometimes overlapping) patterns of innervation of LPBN, providing another potential anatomical driver of our functional maps.

Genetically defined LPBN subpopulations are also spatially organized, with many subpopulations confined to specific subregions of LPBN^22,23^. While we have not examined how our functional groups map onto genetic subtypes, future work could tackle this question using post-hoc genetic identification of imaged neurons as was done previously for the PVH^90^. However, we hypothesize that currently identified genetically defined populations may encompass multiple functional subtypes. For example, LPBN CGRP neurons, which respond to various body signals, were shown to be composed of at least three subtypes based on downstream projection patterns^21^ and to form at least two molecularly distinct subtypes^23^. Consistent with these studies, our fiber photometry recordings showed that CGRP neurons have responses characteristic of both Early and Late Ensure neurons, and our GLM analysis showed mouse-to-mouse variability in how much licking and/or running contributed to CGRP activity **(Figure S8)**. These results suggest that we may have recorded from different CGRP subtypes in different mice, potentially due to small differences in recording location.

Our identification of spatial maps could not have been resolved without our approach employing an angled 1-mm GRIN lens imaging positioned just above LPBN. This method allowed us to access a much larger portion of LPBN in a single mouse with less damage to LPBN as compared to previous imaging studies using lower-profile GRIN lenses inserted directly into LPBN^13,20,24,91,92^. However, our method suffers from a few limitations. First, to verify which LPBN subregions we imaged from and to combine spatial data across mice, we relied on relatively crude mapping of our FOVs to Allen Brain Atlas coordinates. While we believe that we were imaging from PBNdl and PBNel (consistent with the localization of CGRP neurons in **Figure 2C**), we cannot be certain of the precise imaging coordinates within these regions. Second, the surgery to implant the lens just above the PBN was technically very challenging with a low success rate (< 25%), leading to a low throughput despite the large number of neurons recorded across many sessions in successful experiments.

### What does InsCtx input convey to LPBN?

The InsCtx, which encompasses the primary viscerosensory cortex, sends direct feedback to LPBN. However, the function of this input is largely unknown. We found that optogenetic stimulation of InsCtx^LPBN^ axons drives strong activity in the subset of LPBN neurons that is also driven by LiCl and stomach stretch, including CGRP neurons **(Figures 7 and S7G-I)**. InsCtx not only responds to interoceptive stimuli, such as stomach stretch, malaise, and changes in heart and respiratory rate^69,93,94^, but also mediates responses to environmental cues that predict deviations from bodily homeostasis^95–98^. We hypothesize that by activating specific groups of LPBN neurons, InsCtx inputs to LPBN may *simulate* anticipated visceral changes. Consistent with this hypothesis, a recent study showed that specific input from posterior InsCtx to CGRP neurons mediates threat conditioning and can induce anxiety-like behaviors^99^. Through CGRP neurons, InsCtx may induce fear-associated body changes to drive learning of the threat stimulus. Similarly, both InsCtx and LPBN (CGRP neurons in particular) are necessary for conditioned taste aversion (CTA) learning, whereby a novel palatable taste becomes aversive after it is paired with visceral malaise, such as LiCl injection^91,100–102^. We propose that after CTA and upon re-exposure to the conditioned taste, InsCtx^LPBN^ axons may contribute to the activation of LiCl-driven LPBN neurons to simulate malaise and drive avoidance of the previously palatable taste.

We were surprised that input from such a broad region of cortex activated a specific, spatially restricted subset of LPBN neurons. Yet, histological examination also suggested stereotyped, non-uniform InsCtx innervation of LPBN across mice **(Figure 7B)**. This restricted activation was not due to stimulating only a subset of InsCtx^LPBN^ axons, as we observed broad expression of ChrimsonR across InsCtx **(Figure S7A)**. It is also not due to a bias in our FOVs to certain LPBN subregions, as we have observed similar InsCtx^LPBN^-induced focal activation of LPBN neurons during *in vitro* brain slice imaging experiments that include the entire LPBN (data not shown). We suggest that by activating specific subsets of LPBN neurons that encode diverse body states and stimuli (including CGRP neurons; **Figure S7G-I)**, InsCtx can predict various homeostatic deviations (as discussed above) and prime multiple downstream circuits to respond to the expected visceral changes.

This combined optogenetic and imaging strategy highlights a general experimental approach that could be used to study any of the various inputs to LPBN. This method would reveal whether different sources of input to LPBN engage distinct functional groups, enhancing our characterization of these groups as well as our understanding of the roles of specific long-range inputs. Given the distinct spatial localization of axons from different brain regions in LPBN, we suspect that different inputs will converge on different specialized groups of LPBN neurons. Ultimately, this approach could help build a map of LPBN that combines genetic subtype with functional characterization based on responses to both natural and artificial stimuli.

Overall, our work establishes a platform for studying interoception during behavior by combining large-scale neural imaging with monitoring of internal signals and state. Future work could apply this approach to processes and brain regions beyond food intake and LPBN to provide a comprehensive view of the neural circuits underlying body-to-brain communication.

## Acknowledgements

We thank S. Zhang, K. Evans, C. Massengill, J.S. Alvarado, and members of the Andermann lab for helpful discussion and feedback. We thank J. Mathai for use of his LabScope for fluoroscopy experiments. We thank J. Fernando, P. Prasad, D. Guarino, J. DeBolt, Z. Stolberg, and A. Pinilla for animal husbandry and other technical assistance. Authors were supported by NSF GRFP DGE1745303 and NIH F31DC020631 (R.A.E.); NIH F32DK135247 and NIH 5T32DK00751 (K.R.); BBRF Young Investigator Grant (H.K.); The Research Council of Norway Grant 250259 (S.G. and K.L.); Pew Innovation Fund, McKnight Foundation, and Klarman Family Foundation (M.L.A.).

## Author contributions

R.A.E. and M.L.A. conceived of the project. R.A.E., K.R., and M.L.A. wrote the manuscript. R.A.E. and K.R. performed two-photon imaging experiments and data analyses with help from M.L.A. R.A.E. and H.C. performed surgeries, photometry, and brain histology. H.C. and O.A. performed alignment and 3D rendering of coronal sections in the Allen Mouse Brain Common Coordinate Framework. R.A.E., K.R., and H.C. performed X-ray videofluoroscopy experiments, and R.A.E., K.R., H.C., and H.K. analyzed fluoroscopy data with help from M.L.A. T.E.L. provided advice and protocols for X-ray videofluoroscopy. S.G. and K.L. developed the ribo-GCaMP8s sensor. J.E. labeled data for CellPose.

## Methods

## RESOURCE AVAILABILITY

### Lead contact

Further information regarding resources or reagents should be directed to and will be fulfilled by the lead contact, Mark L. Andermann.

### Materials availability

This study did not generate new unique reagents.

### Data and code availability

All photometry and two-photon calcium imaging data and original code reported in this paper have been deposited at: https://github.com/kmruda/PBN_sensing_body_signals

Any additional information required to reanalyze the data is available upon request from the lead contact, Mark L. Andermann.

## EXPERIMENTAL MODEL AND SUBJECT DETAILS

### Animals

All animal care and experimental procedures were approved by the Beth Israel Deaconess Medical Center Institutional Animal Care and Use Committee. Mice were housed in a 12 hour light/dark cycle environment (all experiments were performed during the light cycle) and provided with standard mouse chow (Teklad F6 Rodent Diet 8664; Harlan Teklad) and water a*d libitum*, unless otherwise specified. We used male and female C57Bl/6 or heterozygous *Calca-Cre* mice (age: 8-15 weeks at time of surgery). *Calca-Cre* mice were maintained on a C57Bl/6 background. We used 9 mice (5 male, 4 female) for LPBN two-photon calcium imaging, 6 mice (3 male, 3 female) for CGRP fiber photometry, and 4 mice for videofluoroscopy experiments (2 male, 2 female). Mice were singly housed following surgery and during food restriction.

## METHOD DETAILS

### Surgical procedures

#### Stereotaxic injections

Mice were anesthetized with isoflurane mixed in 100% O_2_ (induction: 3%, maintenance: 1%–2%), and placed into a stereotaxic apparatus (Kopf model 963). Mice were placed on a heating pad (CWE) to maintain their body temperature throughout the surgery. After exposing the skull via a small incision, a small hole was drilled for injection. A pulled-glass pipette with 20–40 mm tip diameter was inserted into the brain, and virus was injected using an air pressure system (Picospritzer; Parker) at a rate of 10-100 nl/min. After five minutes, the pipette was slowly removed for adequate absorption and spread of the virus. For postoperative care, mice were injected subcutaneously with slow-release meloxicam (0.5 mg/kg; ZooPharm). Mice were 8-15 weeks old at the time of injection and given at least one week to recover from surgery before habituation.

We used the following volumes of virus and injection coordinates: InsCtx (200 nl, Bregma: AP: 0.0 mm, DV: −4.1 mm, ML: −4.0 to −4.1 mm), LPBN (200 nl, Bregma: AP: −5 to −5.1 mm, DV: - 3.78 mm, ML: −1.5 to −1.6 mm).

#### Gradient index (GRIN) lens implantation

Mice were stereotaxically injected with AAV-PHP.eB-hSyn-Ribo-jGCaMP8s or AAV-PHP.eB-hSyn-RiboL1-jGCaMP8s into the left LPBN (Bregma: AP: −5.0 mm, DV: −3.78 mm, ML: −1.5 mm) as described above. For experiments involving optogenetic stimulation of InsCtx^LPBN^ axons, AAV1-Syn-ChrimsonR-tdTomato was also injected into left InsCtx (Bregma: AP: 0.0 mm, DV: - 4.1 mm, ML: −4.0 to −4.1 mm). We used a custom-made holder connected to the stereotaxic arms to place a titanium headpost on the skull, tilted at a 40° ML angle, and fixed the headpost to the skull with C&B Metabond (Parkell). We performed a 1.0 mm craniotomy of the skull area at a 40° ML angle relative to LPBN (Bregma: AP: −5.34 mm, ML: −3.4 mm) and aspirated ∼2.0 mm of cerebellar tissue to allow for clean insertion of the GRIN lens. We lowered a 1.0 mm diameter singlet GRIN lens (GRINtech, NEM-100-25-10-860-S-0.5P; 1.0 mm diameter; 4.38 mm length) at a 40° ML angle into the craniotomy to a depth of 2.0 mm from the skull surface (to sit at the surface of LPBN). The lens was secured to the skull first with Vetbond glue (3M) and then with C&B Metabond (Parkell). After completion of the surgery, we protected the top face of the GRIN lens by securing a cut-off tip of an Eppendorf tube (Fisher) around the GRIN lens with Kwik-Cast (WPI). Mice were allowed at least three weeks for recovery before habituation and imaging began.

Following recovery, we prepared the mice for two-photon imaging by cementing a 3D printed plastic funnel onto the headpost, which allowed for the securing of a light shield between the head and objective during imaging.

### Optic fiber implantation

*Calca-Cre* mice were stereotaxically injected with AAV1-Syn-FLEX-GCaMP6s into the left LPBN and with AAV1-Syn-ChrimsonR-tdTomato into the left InsCtx as described above. Next, we performed a 0.5 mm circular craniotomy above LPBN (Bregma: AP: −5.1 mm; ML: −1.6 mm) and implanted an optic fiber (400 µm diameter core; 5.0 mm length; 0.48 numerical aperture; MMFC_400/430-0.48_5.0mm_MF1.25_FLT; Doric Lenses) above LPBN at a depth of −3.6 mm relative to Bregma. The optic fiber and a titanium headpost were fixed to the skull using C&B Metabond (Parkell). Mice were allowed at least one week to recover before habituation began.

### Brain tissue preparation and immunohistochemistry

Mice were terminally anesthetized with tribromoethanol (Sigma Aldrich) and transcardially perfused with 0.1 M phosphate-buffered saline (PBS, pH 7.4) followed by 10% neutral buffered formalin (NBF, Fisher Scientific). To preserve the GRIN lens or optic fiber tract, perfused heads were stored in NBF for at least one day prior to extracting the brain. After brain and implant extraction, the brains were post-fixed overnight at 4°C in NBF and cryoprotected in 20% sucrose. Brains were sectioned coronally on a freezing sliding microtome (Leica Biosystems) at 60 µM thickness and sections spanning LPBN or InsCtx were mounted on coated glass slides (Fisher Scientific) and covered with DAPI-containing Vectashield Anti-fade Medium (Vector Laboratories). Coverslips (Fisher Scientific) were placed over the sections and sealed with nail polish. Fluorescent images were captured at 10x magnification with an Olympus VS200 ASW slide scanner microscope.

### Alignment of brain slices to Allen Brain Atlas Common Coordinate Framework

Brightfield images of the brain slices containing LPBN were captured with an Olympus VS200 ASW slide scanner microscope. We used the program QuickNII^103^ (NITRC) to manually align individual brain slices to the Allen Mouse Brain reference atlas using visible and distinct anatomical structures within each slice (e.g., shape of the superior cerebellar peduncle tract, *scp*). In parallel, we used ImageJ to create binary masks of the GRIN lens tract in each LPBN-containing brain slice for reconstruction of GRIN lens placement throughout the brain **(Figure S2A)**. Also using ImageJ, we created binary masks of the location of each two-photon FOV within the 1 mm GRIN lens (one FOV per mouse) by aligning vasculature in the two-photon FOV to that in epifluorescence images of the brain area seen through the whole GRIN lens. Next, we used Brainrender^104^ to reconstruct the GRIN lens and FOV locations in 3D Allen Brain Atlas Common Coordinate space based on the GRIN lens tract and FOV masks, and the reference atlas-aligned slices. This process was completed on the brains from all mice used in imaging experiments, allowing for comparison of GRIN lens placement as well as merging of spatial maps across mice as in **Figure 5**.

### Mouse behavior training

Mice underwent a systematic habituation process to mitigate any stress from head fixation. Initially, over 2-3 days, mice were incrementally introduced to head fixation while allowed to run freely on a 3D-printed running wheel. If the mice displayed any signs of stress, they were immediately removed, and additional head-fixation sessions were added until there were no visible signs of stress. Subsequently, mice were food-restricted to 85-90% of their free-feeding body weight (∼2.5 g chow/day) and were trained to lick for Vanilla Ensure (a high-calorie liquid meal replacement; Abbott) via hand feeding with a syringe. Once mice were habituated to hand feeding, we trained mice to lick for one drop of Ensure (22 µl) every 20 s (as in our imaging experiments) using MonkeyLogic^105^ (NIMH/NIH) for automatic Ensure delivery. Mice were trained for 30 min – 1 hour per day until they learned to lick for each Ensure delivery and mostly withhold licking during the 20 s ITI period. Prior to the free-feeding experiments **(Figure 6)**, we additionally trained mice to associate licking the lickspout with automatic delivery of Ensure (10 µl drops with a minimum ITI of 2 s between deliveries).

### Two-photon calcium imaging with interoceptive stimulus delivery

We performed two-photon calcium imaging using a resonant-scanning two-photon microscope (Neurolabware) at 15.63 frames s^-1^ and 796 x 512 pixels/frame. We used an InSight X3 laser (Spectra-Physics) tuned to either 920 or 960 nm for GCaMP fluorescence excitation and performed imaging with a 10x 0.5 NA air objective (TL10X-2P; ThorLabs). Our imaging fields of view were at a depth of 100-300 μm below the face of the GRIN lens and targeted to the surface of LPBN. We imaged 1-2 planes per mouse, spaced at least 30 µm apart. Laser power for excitation ranged from 20-60 mW at the front aperture of the objective; however, power at the brain was much less due to power loss through the GRIN lens.

### Ensure, tail shock, LiCl, and CCK sessions

Imaging sessions were ∼80 minutes long and consisted of the following runs: (1) 5 minute baseline period (no stimuli presented), (2) 70 randomly interleaved Pavlovian Ensure (22 µl drops), tail shock (0.3 mA; 2 x 50 ms pulses; 100 ms inter-shock interval), and blank trials with a 20 s ITI in a 4 Ensure: 2 tail shock: 1 blank trial ratio (40 Ensure, 20 tail shock, and 10 blank trials total in the run), and (3) a 5 minute baseline period followed by an i.p. saline injection, 10 minute post-injection period, i.p. LiCl or CCK injection, and a 25 minute post-injection period. Ensure was delivered to the head-fixed mice via a syringe pump (Harvard Apparatus) connected to a custom 3D-printed lickspout. Tail shocks were delivered via two electrode pads (Covidien; Series S) that were wrapped around the base of the mouse’s tail, and current (0.3 mA) was delivered to the electrode pads using a stimulus isolator (Iso-Flex; AMPI). For runs involving pharmacological injections, we first injected 350 µl of sterile 0.9% saline as a control. After 10 minutes, we injected 350 µl of either 0.2M LiCl (14 ml/kg) or CCK (10 µg/kg) and recorded neural activity for an additional 25 minutes. Sterile 0.9% saline was stored at room temperature (25°C before use). LiCl was prepared in sterile dH_2_O and stored at 4°C before use. CCK was freshly prepared in sterile 0.9% saline before injection. Mice were extensively habituated to needle injections prior to recordings.

### Optogenetic InsCtx^LPBN^ axon stimulation

In a subset of mice (n = 5) expressing ChrimsonR-tdTomato in InsCtx^LPBN^ axons, we added optogenetic stimulation trials to the imaging sessions described above. These sessions consisted of the same runs as above, except for the second run, which instead had 90 randomly interleaved Ensure, tail shock, InsCtx^LPBN^ optogenetic stimulation (2 s; 2.5 mW; 15.63 Hz; 8 ms pulses), and blank trials in a 4 Ensure: 2 shock: 2 optogenetic stimulation: 1 blank trial ratio (40 Ensure, 20 shock, 20 optogenetic stimulation, and 10 blank trials total). For optogenetic stimulation of ChrimsonR, light was delivered through the imaging objective and GRIN lens using a 617 nm LED (M617L3; ThorLabs) controlled by an LED driver (T-Cube, ThorLabs) that was synchronized with imaging acquisition (as in Lutas et al. 2022^106^ and Reggiani et al. 2023^107^). To protect the PMT (H11706-40; Hamamatsu) from optogenetic stimulation light, the PMT was gated for 10 ms at the onset of stimulation. This gating resulted in the top 16% of each frame being blank during photostimulation (2 s).

### Ensure free feeding sessions

Mice were given 30 minutes of free access to Ensure during imaging. Ensure delivery (10 µl) was contingent on licking and mice could receive a maximum of 1 drop every 2 s. These Ensure delivery parameters caused the mice to lick continuously for Ensure until they were sated (usually after about 2-3 mL of consumption). To ensure that mice continuously licked, we trained them on this feeding paradigm for at least 3 days (1 hour/day) prior to imaging.

### Anesthetized sessions with stomach balloons

The day after performing an awake imaging session with Ensure, tail shock, and LiCl delivery, we recorded anesthetized responses to gastric stretch. We anesthetized mice with isoflurane (3% induction, 0.5 - 1.5% maintenance) and implanted a latex stomach balloon (Harvard Apparatus, 73-3479) attached to an oral gavage needle (Cadence Science, 9920) via surgical sutures into the forestomach (as in Ran et al. 2022^28^). The abdomen was closed with sutures and surgical glue (Vetbond, 3M). Mice were then carefully transported to the two-photon imaging setup for recording, with a heating pad in place of the running wheel. Heart rate, breathing, body temperature, and toe pinch responses were monitored closely during these sessions. Anesthetized recordings consisted of two runs: first, a baseline run, and second, a stomach inflation run. In this second run, the stomach was inflated to a volume of 0.45 mL in 3 s (at a rate of 9 mL/min), held at 0.45 mL for 14 s, deflated over 3 s, and held at 0 mL for 14 s. These trials were repeated up to 32 times. In a subset of mice, we also included runs involving mild tail shock (same as in awake sessions except at 0.4 mA) and/or InsCtx optogenetic stimulation (same as in awake sessions). Mice remained under anesthesia during these sessions and were euthanized immediately afterwards.

### Fiber photometry with interoceptive stimulus delivery

Fiber photometry recordings were performed in head-fixed mice on a running wheel that were extensively habituated to head fixation and Ensure consumption as described above. We coupled fiber optic cables (1 m long; 400 µm core; 0.48 NA; Doric Lenses) to implanted optic fibers with zirconia sleeves (Precision Fiber Products) and placed black heat shrink material around the fiber coupling to block external sources of light from interfering with the recordings. The black heat shrink material also blocked the mice from seeing the pulses of red light used during optogenetic stimulation. We used the same fiber optic cable to collect GCaMP fluorescence signals (excited by a 465 nm Plexon LED and driver, 30-100 µW, sampled at 1 kHz) and to deliver light for optogenetic stimulation (620 nm Plexon LED and driver, 2.5 mW, 20 Hz, 10 ms pulses). Excitation light (for both photometry and optogenetics) and emission light were passed through a fluorescence minicube (FMC4_E(460-490)_F(500-550)_O(580-650)_S; Doric Lenses), and emission light was collected by a photoreceiver (Newport 2151), which was then demodulated by a lock-in amplifier (SR830; Stanford Instruments). We used a NI-DAQ (PCIe-6321; National Instruments) and custom MATLAB scripts (MathWorks) to collect digitized photometry signals. We performed stimulus presentations (Ensure, tail-shock, InsCtx^LPBN^ optogenetic stimulation, stomach stretch, LiCl injection, and saline injection) in the same manner as described above for the two-photon imaging experiments.

### Behavioral tracking during two-photon imaging and fiber photometry recordings

For pupil tracking, we acquired videography of the eye using a GigE Vision camera (Dalsa) with a 60 mm lens (Nikon MicroNikkor). During two-photon imaging, the pupil was illuminated by infrared laser light exiting out through the pupil, and during fiber photometry, we used an LED array to illuminate the pupil. We recorded running speed in MATLAB using a custom-made encoder (Arduino) coupled to the 3D-printed running wheel. We tracked licking behavior via a capacitance-sensing lickspout (3D printed in a metal-containing filament connected to the MPR121 capacitance sensor). See below for details of analyses of behavioral variables.

### X-ray videofluoroscopy in head-fixed mice

We adapted previous methods for X-ray videofluoroscopy^31^ in freely moving, small animals to a head fixation setting in order to track the real-time movement of Ensure through the GI tract. Fluoroscopy recordings were performed with help from John Mathai’s lab at BIDMC/Harvard using a low energy fluoroscope (23 kV, 200 A, and 0.54-mm Al filter LabScope; Glenbrook Technologies, Newark, NJ). Fluoroscopy data was acquired at 30 frames s^-1^ with a voltage in the range of 34.6 - 35.4 kV and a current of 0.216 A. Mice were habituated to head-fixation and body restraint in a custom conical tube and were trained to lick Ensure as above. In half of the fluoroscopy sessions, the stimulus delivery was identical to two-photon imaging sessions, with a mix of mild tail shocks, blank trials, and small drops of Ensure. In the other half of sessions, mice were presented with just Ensure every 20 s. To minimize radiation exposure to mice, sessions were limited to 30 minutes a day for four days total. We recorded licking activity through a lickspout as described above, and synchronized fluoroscopy recordings with licking records based on licking activity readily observed from fluoroscopy.

Ensure was mixed with the oral contrast agent Iohexol (350 mg iodine/ml, Omnipaque, GE HealthCare) for visualization under X-ray imaging. Preliminary tests showed that a ratio of 1 Ensure: 1 Iohexol produced strong contrast in recordings while maintaining similar viscosity and palatability to unmixed Ensure (data not shown). Following food restriction and Ensure lick training (see above), mice readily licked for the Ensure/Iohexol mixture (see **Figure 1C**). We also confirmed that the viscosity of the liquid food does not affect GI tract temporal dynamics by performing the same experiments with Ensure thickened with SimplyThick (data not shown).

## QUANTIFICATION AND STATISTICAL ANALYSIS

### Two-photon imaging analysis

#### Image registration and timecourse extraction

We used Suite2p^108^ to perform image registration of each two-photon calcium imaging FOV. We trained a custom CellPose^109^ model to identify ribo-GCaMP8s-expressing cell body regions of interest (ROIs) from the mean image of each motion corrected FOV, and we extracted fluorescence timecourses by averaging the pixels within each binarized ROI mask (F_ROI_(t)). We calculated neuropil activity as the median value of the surrounding region of each ROI excluding pixels that belonged to any other surrounding ROI (F_neuropil_(t)). We performed neuropil correction on each ROI’s fluorescence timecourse by first subtracting the scaled neuropil timecourse (scaled by 0.7) from the ROI’s fluorescence timecourse and then adding back the median value of the neuropil trace: F_neuropil_corrected_(t) = (F_ROI_(t) - 0.7*F_neuropil_(t)) + F_median_neuropil_. We then Z-scored each ROI’s neuropil corrected timecourse across all runs within a single imaging session.

### Alignment of cell masks across days

During acquisition, care was taken to image FOVs that were as similar as possible across days. We then used CellReg^110^, which computes a probabilistic model of spatial ROI patterns to find the same neurons across two days of recordings. The spatial footprint of matched neurons was visually verified, and we observed that many response properties were stable across days of recordings **(Figures 2E and S2E)**.

### Analysis of behavioral variables

For pupil area, we used a custom trained DeepLapCut^38^ model to track the outline of the pupil with 8 points in each recorded video frame. To calculate and extract pupil area traces, we used the 8 outline points to fit an ellipse around the pupil in each frame and then calculated the area of the ellipse using the ellipse area formula (π x r_major axis_ x r_minor axis_). We then smoothed the pupil area traces with a 1-s running median filter. Finally, we normalized the pupil area of each frame to the maximum pupil area from a given imaging session, such that pupil area values would be between 0 and 1 (Pupil_area_normalized_(t) = Pupil_area(t) / Pupil_area_maximum_).

For licking, we calculated lick rate (licks/s) using a 1-s sliding window. For running, we extracted running speed traces (in cm/s) from the encoder attached to the wheel and smoothed the traces with a 0.33-s moving mean. We aligned pupil, licking, and running traces to the onset of each stimulus as described below for neural activity.

### Single neuron stimulus selectivity

For stimuli that were repeated multiple times in a session (Ensure, tail shock, running, stomach stretch, and InsCtx^LPBN^ optogenetic stimulation) we determined stimulus-evoked responses of each neuron by aligning each neuron’s Z-scored fluorescence trace to the onset times of each stimulus (or first lick after Ensure delivery for Ensure) and considering activity in the 2 s prior to stimulus onset up to the 15 s after stimulus onset. For heatmap visualizations, we normalized the evoked response of each neuron on each trial to each 2-s baseline period (by subtracting the mean of this baseline period) and took the mean of each neuron’s normalized trace across all stimulus trials (a 5-s baseline period was used for anesthetized stomach stretch trials). For the i.p. injection stimuli (saline, LiCl, and CCK), we considered activity in the 5 minutes prior to the injection time up to either 10 min (for saline) or 25 min (for LiCl and CCK) after the injection and normalized to the baseline period as above. For heatmap visualizations of injection responses, we additionally smoothed each neuron’s trace using the mean values from a 10-s moving window. To find running bouts, we first limited analyses to baseline runs, ITI periods 6 s to 19 s after stimulus delivery during stimulus runs, and any post-stimulus runs. Running bouts were defined as 1 s of no running (running speed < 1 cm/s for all bins, which were 128 ms wide) followed by at least 1 s of running (running speed ≥ 1 cm/s for all bins).

To categorize neurons as driven by a stimulus, we used different procedures for the repeated stimuli and the injection stimuli because the injection stimuli were only delivered once per session. For Ensure, tail shock, stomach stretch, and InsCtx^LPBN^ stimulation, we first tested whether a neuron’s response was statistically significant, and then we categorized the response as activated or suppressed based on a Z-score threshold. We compared activity in the 2-s baseline period (or 5 s for stomach stretch) to activity up to 10 s (for Ensure, shock, and stomach stretch) or 2 s (for InsCtx^LPBN^ opto stim) after the stimulus. We binned the post stimulus activity by taking the mean of every 250 ms and then performed a Wilcoxon Signed-Rank test, followed by a Bonferroni correction for multiple comparisons, on each 250 ms bin compared to the mean of the baseline activity. We considered a response as significant if 2 or more consecutive bins were statistically significant (P < 0.05). We then took the maximum and minimum response (Max_activity_, Min_activity_) of a neuron’s mean stimulus-evoked activity in the post-stimulus windows listed above and categorized the neuron as activated if Max_activity_ ≥ 0.5 Z and suppressed if Min_activity_ ≤ −0.5 Z. For anesthetized stomach stretch recordings, we used a slightly more permissive criterion because of reduced neural activity in the anesthetized state: responses were considered activated if Max_activity ≤_ 0.5 Z or if the average activity in the 10 s after stomach stretch expansion (Avg_activity_) > 0 Z. Neurons driven by running were defined as those having a fractional deviance explained by running (from GLM analyses, see below) > 0.65.

For the i.p. injection stimuli (saline, LiCl, CCK), we calculated an auROC value for each neuron by comparing activity in the 5 min baseline period to activity in the 10 min (saline) or 20 min (LiCl, CCK) post injection period. auROC values have a range of 0-1: a value of 0 indicates a clear decrease in activity relative to the baseline, a value of 1 indicates a clear increase in activity relative to the baseline, and a value of 0.5 indicates no difference in activity between the baseline and post-injection period. We categorized a neuron as activated by the injection if auROC ≥ 0.6, suppressed if auROC ≤ 0.4, and not driven for all other auROC values.

### Categorizing Ensure responses

For neurons that were significantly activated by Ensure consumption, we categorized their responses as Early (transient or sustained) or Late based on response onset time and duration. First, we smoothed each neuron’s mean response to Ensure with a 0.33 s moving mean. Next, we defined onset time as the time to reach 50% of the peak smoothed activity relative to the first lick. For response duration, we also estimated the fall time, which we defined as the first-time post-stimulus activity returned to <50% of peak activity after the peak response. We then calculated response duration as: fall time – onset time. We plotted the distribution of onset times **(Figures 1G and S1E)** and response durations **(Figure S1E)** to set thresholds for our different groups of neurons. Early Ensure neurons: onset time < 0.9 s (transient: response duration ≤ 0.9 s; sustained: response duration > 0.9 s); Late Ensure neurons: onset time ≥ 0.9 s.

### Changes in Ensure trial-evoked activity

To determine whether a neuron’s Ensure trial-evoked activity changes over a session, we found the peak activity of each trial by averaging the top 4 activity values in the 15 s post-stimulus period of each trial. For the heatmap visualization in **Figure 1F**, we took the mean of the peak activity from every 4 trials and plotted those values for all Ensure-activated neurons. We then used a paired t-test to compare the peak activity values from the last 10 trials to those from the first 10 trials. We performed this procedure for each neuron on two consecutive days of identical imaging sessions, and we considered a neuron as having a significant change in Ensure-evoked activity if its change was significant (P < 0.05) and in the same direction (increase or decrease) on both days. We calculated the direction of the change from (mean of peak activity_Last 10 trials_ – mean of peak activity_First 10 trials_). Positive values indicated an increase in trial-evoked activity and negative values indicated a decrease.

### Slow changes in ongoing activity

We defined a neuron’s ongoing activity as activity unrelated to any stimulus and outside of active licking and running bouts (“quiet waking” activity). For baseline runs, we removed activity during periods where running ≥ 1 cm/s and licking ≥ 1 lick/s. For Ensure trial runs, we concatenated activity from the last 10 s of each 20 s ITI following an Ensure trial, removing periods with running ≥ 1 cm/s and licking ≥ 1 lick/s. The free feeding runs did not have built in ITIs so we removed activity during running bouts ≥ 1 cm/s and during lick bouts > 0.5 licks/s. We used a lower lick threshold here to ensure that included activity was not within active consumption bouts – we only included activity during periods where single licks were at least 2 s apart.

We found neurons with significant ongoing activity changes during an hour-long imaging session by comparing the ongoing activity in the last 2 minutes of the session (“End” activity) to that in the first 2 minutes of the session (“Beginning” activity). We binned the beginning and end activity by taking the mean of every 5 s and then performed a paired t-test to determine which neurons’ activity was significantly different at the end of the session compared to the beginning of the session (P < 0.05). We followed this procedure for each neuron on two consecutive days of imaging, and we considered a neuron as having a significant change in ongoing activity if the change was significant on both days of imaging. We also calculated the change in ongoing activity of each neuron as the mean activity in the last two minutes minus the mean activity in the first two minutes.

### Generalized linear model (GLM)

We quantified the contribution of behavioral variables to each neuron’s activity by fitting a Gaussian GLM using the GLMnet package^111,112^. Each behavior trace (licking rate, running speed, pupil area, and change in pupil area) was convolved with a Gaussian kernel (1 s wide) to create a basis function. This was repeated for each trace for delays from −3 to 8 s (with 0.5 s spacing) to account for various delays between behavioral variables and neural activity. We did not include Ensure or tail shock delivery as input variables because their influence on neural activity became split between stimulus delivery and other behavioral variables (for example, coefficient weights for a lick-driven neuron would be unreliably split between licking rate and Ensure delivery). For neural activity, we first calculated the fractional change in calcium fluorescence using ΔF/F(t) = (F_neuropil_corrected_(t) - F_0_(t)) / F_0_(t), where F_0_(t) is the 10th percentile of a 30 s sliding window of F_neuropil_corrected_(t) (as in Livneh et al. 2020^60^). We then Z-scored ΔF/F traces for comparison across mice and recording days. The GLMs were trained on 75% of the data, and all activity predictions and model performances reported are from the remaining 25% testing set. These test and train sections were interleaved in 4-6 blocks throughout recordings to account for any slow (minutes-long) changes in neural activity. Model performance was reported using fractional deviance between the predicted and actual neural activity. We used α = 0.1 for elastic net regularization.

The GLM with all behavioral variables included is referred to as the “full” model. To determine the extent to which each behavioral variable was sufficient to explain neural activity, we repeated the above process, using only one behavior trace at a time. The fractional deviance is the deviance explained by one behavioral variable divided by the deviance explained under the full model. Because some behavioral variables are highly correlated with each other (for instance, running and pupil area; **Figure 4D**), their fractional deviances are not mutually exclusive. We note, however, that running and licking are minimally correlated **(Figure 4)**, and so their fractional deviance contributions to neural activity are not likely to be redundant.

### Spatial clustering

We quantified the spatial organization of functional responses with the following procedure: for each neuron in a FOV, we considered an annulus a certain distance away from the neuron’s center location. We computed the fraction of neurons with centroids within that annulus area that were in the same stimulus response category (defined below). This was repeated for annulus distances from 0 to 1000 µm (with a step size and thickness of 25 µm), and then repeated and averaged across all neurons in the group with the same response category to yield probability plots in **Figures 5C-D and S5G-H**. Shuffled probability curves were found by the same procedure with randomly shuffling the stimulus response category of neurons and averaging over 100 shuffle iterations. **Figures 5B and S5E** show clustering probability curves that are averaged over all mice (and expressed as the number of standard deviations away from the average shuffled probability). FOVs with less than 5 neurons in a response category were not included in the analysis. We determined the significance of these clustering curves by averaging the probability between 0-100 µm and comparing to the shuffled values with a Wilcoxon rank sum test. P values in mice with more than one imaged plane were Bonferroni corrected. For Ensure response clustering, we considered the spatial organization of Late vs. Early Ensure neurons and did not include neurons that were not activated by Ensure. For all other stimuli (LiCl, running, and stomach stretch), the two response categories were: (i) neurons activated by the stimulus vs. (ii) the other neurons not activated by the stimulus.

### Fiber photometry analysis

We downsampled fiber photometry traces from 1 kHz to 100 Hz, smoothed the traces using a 250 ms moving average, and Z-scored the traces using all runs in a given day from a given mouse. To determine stimulus responses, we aligned Z-scored photometry traces to the onset of each stimulus trial and normalized the post-stimulus response to the baseline pre-stimulus period. We then took the mean across all trials for individual mice and across all mice where specified. For InsCtx^LPBN^ optogenetic stimulation, we considered activity in a 2 s pre-stimulus baseline period followed by a 15 s post-stimulus period. For LiCl injection, we considered activity in a 5 min pre-stimulus baseline period followed by a 25 min post-stimulus period. For stomach stretch, we considered activity in a 5 s pre-stimulus period followed by a 50 s post-stimulus period (including 3 s inflation, 30 s holding, and 3 s deflation periods). We extracted behavioral variables (running, licking, pupil) and aligned their traces to each stimulus onset as described above for two-photon imaging. GLM analyses were also performed identically as for imaging data, using bulk activity in photometry traces for neural activity instead of individual neurons.

### X-ray videofluoroscopy analysis

To determine the timing of consumption events, we hand-labeled frames to identify the first lick of a trial, each swallow of the Ensure bolus after that first lick, and stomach entry. Stomach exit data **(Figure 1G)** was found from the first seven trials of a recording session for which food entry from stomach to duodenum entry was clearly visible **(Video S2)**. We repeated the visual inspection to select trials manually across three people to verify there was no subjective difference in event tracking. To quantify stomach expansion over the course of a recording, we first trained a DeepLabCut model to track 10 points along the perimeter of the stomach (**Figure 1B** and **Video S4**; this model also included points along the diaphragm, spine, and ventral surface of the mouse as described below). Stomach area was extracted using a spline fit of these perimeter points and reported either as a Z-score **(Figure 1I)** or raw area **(Figure S1D)**. To create kymograph visualizations **(Figure 1F)**, we first corrected for motion artifacts caused by breathing and postural adjustments using BigWarp^113^, an ImageJ plugin implementing Thin Plate Spline deformation^114^. Briefly, anchor points along the spine, torso, jaw and diaphragm were detected by DeepLabCut **(Figure 1B)**. Anchor points from a reference frame were used to register all other frames within a recording session. The resulting deformations were uniformly applied across fluoroscopy frames. Next, we constructed the kymographs from the aligned frames by manually tracing the esophagus from the throat to the stomach entry and fitting a spline to these traced points. We placed 50 equidistant circular ROIs, each with a radius of 10 pixels, along this spline. By calculating the mean pixel intensity within each ROI, we generated an intensity profile along the esophagus across time, consistently applying this method to all frames within each trial.

## KEY RESOURCES TABLE

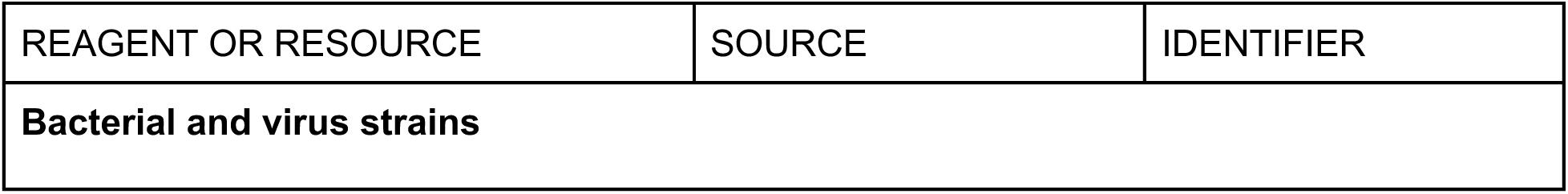
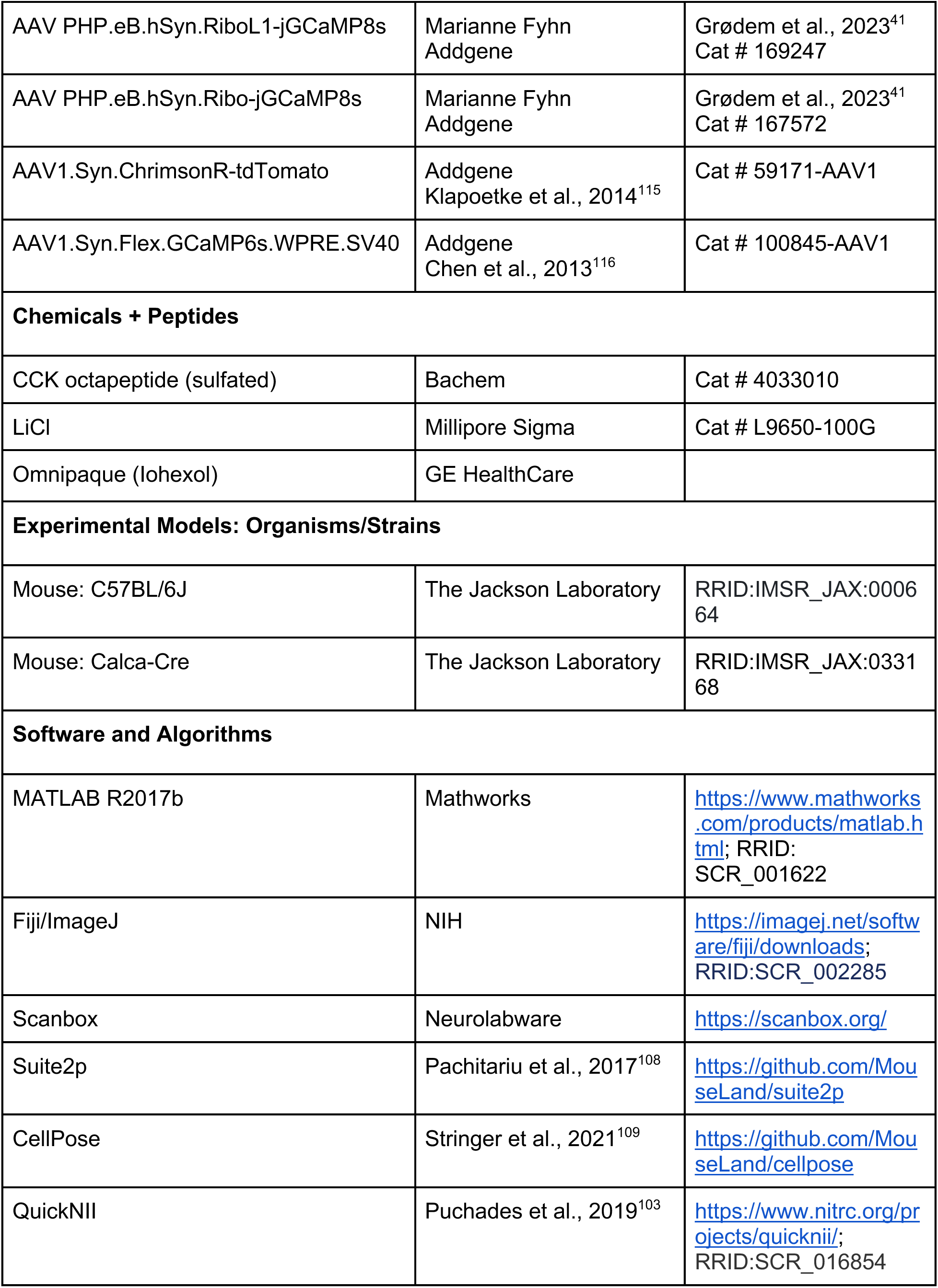
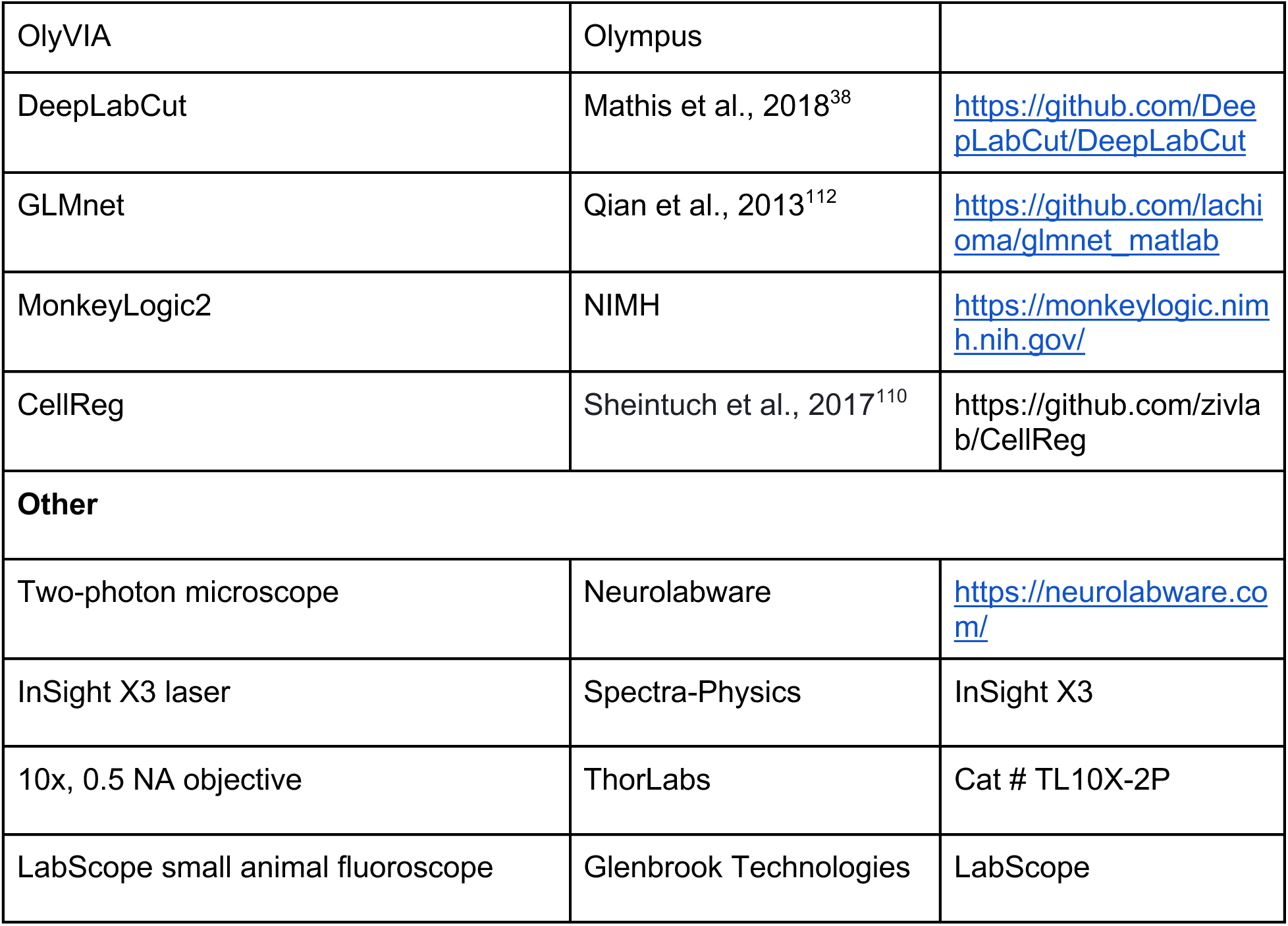

---

**Figure S1, Related to Figure 1:**
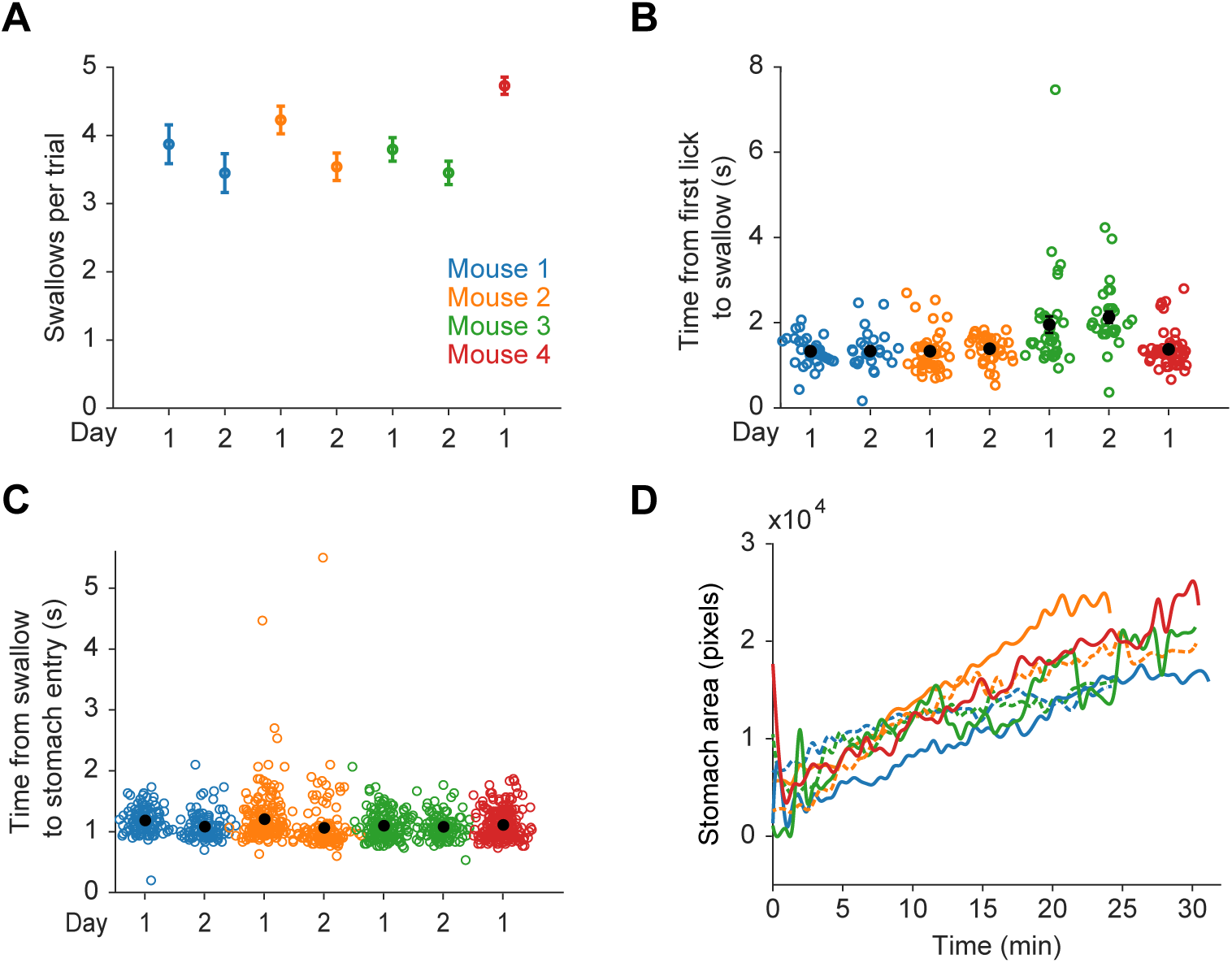
Food transit times and stomach size from individual mice. (A) Average number of swallows per Ensure trial (n = 32-40 trials) for each mouse on two recording days. Error bars are +/-SEM. Mouse color legend is the same as here for B-D. (B) Time from first lick to swallow across all recording sessions. Each colored dot is a single trial, while the black dots are averages from a single mouse and day. Values are not significantly different across days (P = 0.0833, n = 3 mice; Friedman test). Pairwise comparisons between mice were not significant except for Mouse 3, which was significantly different than all the other mice (P << 0.001; Kuskal-Wallis test with Bonferroni correction). (C) Same as (B) for esophageal transit time (from swallow to stomach entry). Values are not significantly different across days (P = 0.0833, n = 3 mice; Friedman test). Pairwise comparisons between mice were not significant except for Mouse 1, which was significantly different than Mouse 2 and Mouse 3 (P = 0.01 and 0.03, respectively; Kuskal-Wallis test with Bonferroni correction). (D) Traces showing stomach area (calculated from DLC labeled points; see Methods) over time on 2 separate recording days for each mouse. Mice ingested ∼1.5 ml of Ensure per session. Solid line: Day 1; Dashed line: Day 2.

**Figure S2, Related to Figure 2.**
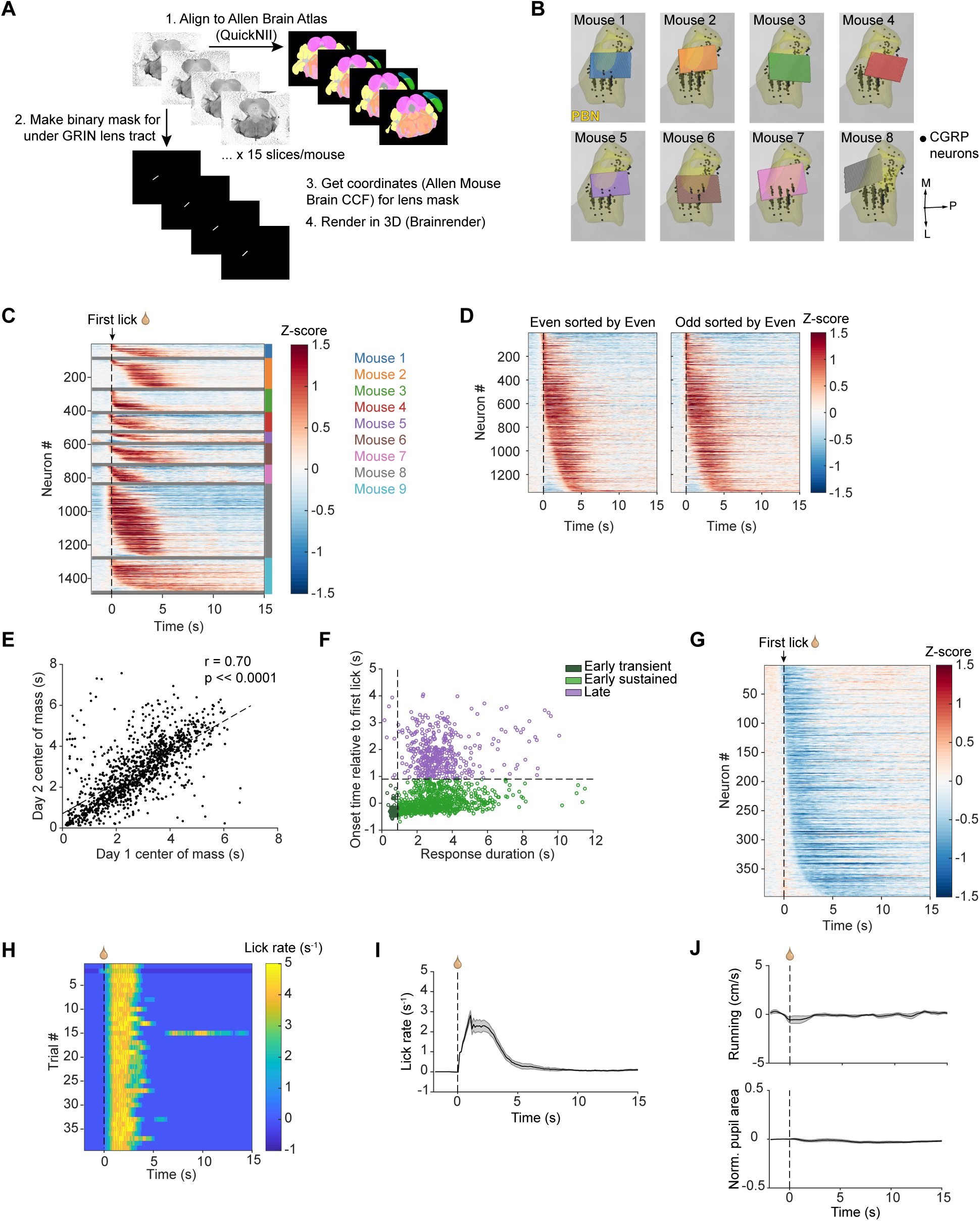
Further analyses of LPBN neural responses to Ensure consumption. (A) Schematic of pipeline for aligning coronal sections to the Allen Mouse Brain Common Coordinate Framework and rendering imaging FOVs in 3D space. (B) 3D rendering of the FOVs from each mouse after alignment to the Allen Mouse Brain Common Coordinate Framework. Gray: 3D outline of mouse brain. Yellow: 3D outline of entire PBN. Black dots: CGRP neurons. P, posterior; M, medial; L, lateral. (C) Heatmaps showing the mean Ensure response of single Ensure-activated neurons from individual mice. Each row is the activity of a single neuron and horizontal gray lines divide the data from each mouse. Neurons are sorted per mouse by Ensure onset time, highlighting that Early and Late Ensure neurons were observed in all mice. (D) Left: Heatmap showing the mean response of single Ensure-activated neurons (n = 1351 neurons from 9 mice) calculated from only the <u>even-numbered trials</u> (n = 20 trials). Right: Heatmap showing the mean response of the same neurons calculated from only the <u>odd-</u> <u>numbered trials</u> (n = 20 trials). In both heatmaps, neurons are sorted by onset time from the mean traces calculated using even numbered trials. (E) Correlation of center of mass of Ensure responses on Day 1 with that on Day 2 for all recorded Ensure-activated neurons. (F) Scatter of Ensure response duration versus response onset time (relative to time of first lick) for all recorded Ensure-activated neurons (n = 1351 neurons from 9 mice), colored by Early transient, Early sustained, and Late groups. Vertical dashed line at 0.9 s indicates boundary between transient and sustained neurons. Horizontal dashed line at 0.9 s indicates boundary between Early and Late neurons. (G) Heatmap showing the mean response (n = 40 trials) of all Ensure-suppressed neurons (n=397 neurons from 9 mice; see Methods), aligned to the first lick and sorted by response onset time. (H) Heatmap showing the lick rate in response to Ensure delivery during individual trials from an example mouse. Each row is the lick rate from a single trial, aligned to the first lick of each trial. (I) Mean lick rate across all Ensure trials from all mice (n = 360 total trials from 9 mice), aligned to first lick per trial. Error bars: +/- SEM. (J) Mean running speed (top) and normalized, baseline-subtracted pupil area (bottom) across all Ensure trials from all mice (n = 360 total trials from 9 mice), aligned to first lick per trial. Error bars: +/- SEM.

**Figure S3, Related to Figure 3:**
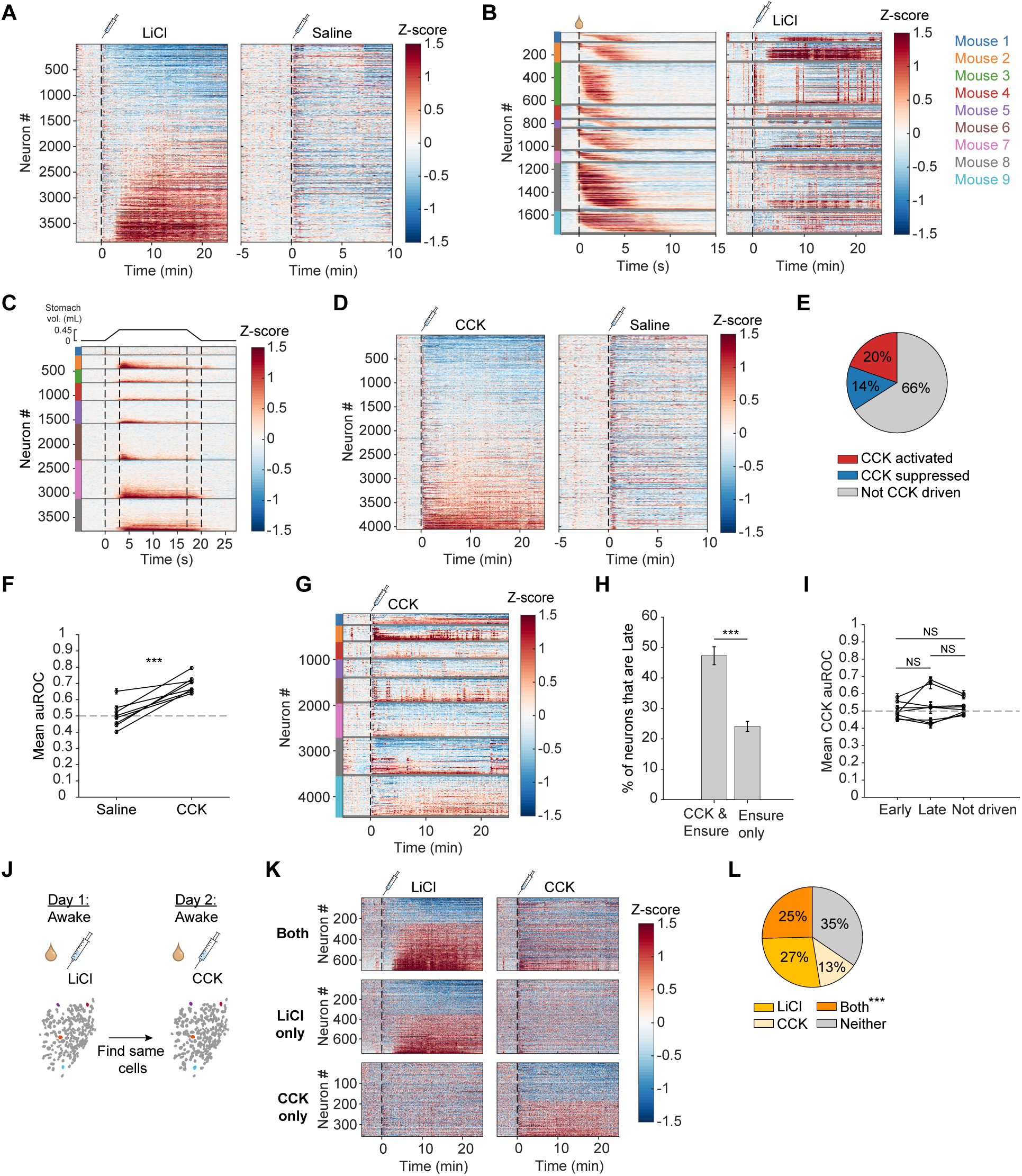
Further analyses of responses to LiCl and stomach stretch; analysis of responses to CCK injection. (A) Left: Heatmap showing the response of single LPBN neurons (n = 3866 neurons from 8 mice) to i.p. LiCl injection. Right: Heatmap showing the response of the same neurons to i.p. injection of 0.9% saline. Neurons in both heatmaps are sorted by LiCl auROC values. (B) Left: Heatmaps showing the mean Ensure response of single Ensure-activated neurons from individual mice. Right: Heatmaps showing the response of the same neurons to LiCl injection. For both Left and Right, each row is the activity from a single neuron and horizontal gray lines divide the data from each mouse. Neurons are sorted per mouse by Ensure onset time. Colors in legend of mouse identity (right) also apply to C and G. (C) Heatmaps showing the response of single LPBN neurons to stomach stretch for individual mice. Each row is the activity from a single neuron and horizontal gray lines divide the data from each mouse. Neurons are sorted per mouse by the mean activity in the first 9 seconds post balloon inflation. (D) Left: Heatmap showing the response of single LPBN neurons (n = 4053 neurons from 8 mice) to i.p. CCK (10 μg/kg) injection. Right: Heatmap showing the response of the same neurons to i.p. injection of 0.9% saline. Neurons in both heatmaps are sorted by CCK auROC values. (E) Pie chart demonstrating the percentage of CCK activated, CCK suppressed, and not CCK driven neurons within the whole population of recorded neurons. CCK activated: 796/4053; CCK suppressed: 584/4053; Not driven: 2673/4053. (F) Mean auROC value of all CCK-activated neurons in each mouse (n = 8 mice) for saline injection and for CCK injection. Dashed line at auROC = 0.5 indicates no response. Each dot represents the mean auROC value from one mouse and values for saline and CCK are connected for a given mouse. ***P = 3.5 x 10^-4^, paired t test. (G) Heatmaps showing the response of single LPBN neurons to CCK injection from individual mice. Each row is the activity from a single neuron and horizontal gray lines divide the data from each mouse. Neurons are sorted per mouse by CCK auROC values. (H) Percent of neurons that is Early or Late onset for the CCK & Ensure activated group (47.3% Late) vs. the group only activated by Ensure (24.1% Late). ***P = 1.3 x 10^-12^, two-proportion Z test. (I) Mean auROC value for CCK injection for Early Ensure neurons, Late Ensure neurons, and neurons not driven by Ensure. Each dot is the mean from an individual mouse and values for a given mouse are connected by a line. Early-Late: P = 1.2; Early-Not Driven: P = 0.49; Late-Not Driven: P = 2.8, paired t test, Bonferroni-corrected post hoc comparisons. (J) Schematic for experiment and analysis to compare LPBN neuron responses to LiCl and CCK. (K) Heatmaps showing responses of single LPBN neurons to LiCl (left) or CCK (right) injection for neurons that are driven by both CCK and LiCl (top row, n = 701), neurons driven only by LiCl (middle row, n = 748), and neurons driven only by CCK (bottom row, n = 357). Data is from 8 mice. (L) Pie chart demonstrating the percentage of neurons driven by both LiCl and CCK, only LiCl, only CCK, or neither LiCl nor CCK. Activated by CCK and LiCl: 701/2758; activated only by CCK: 357/2758; activated only by LiCl: 748/2758; Not driven: 952/2758. ***P << 0.001 for the number of neurons responsive to both LiCl and CCK expected above chance, Fisher’s exact test. Error bars are +/- SEM unless otherwise indicated.

**Figure S4, Related to Figure 4.**
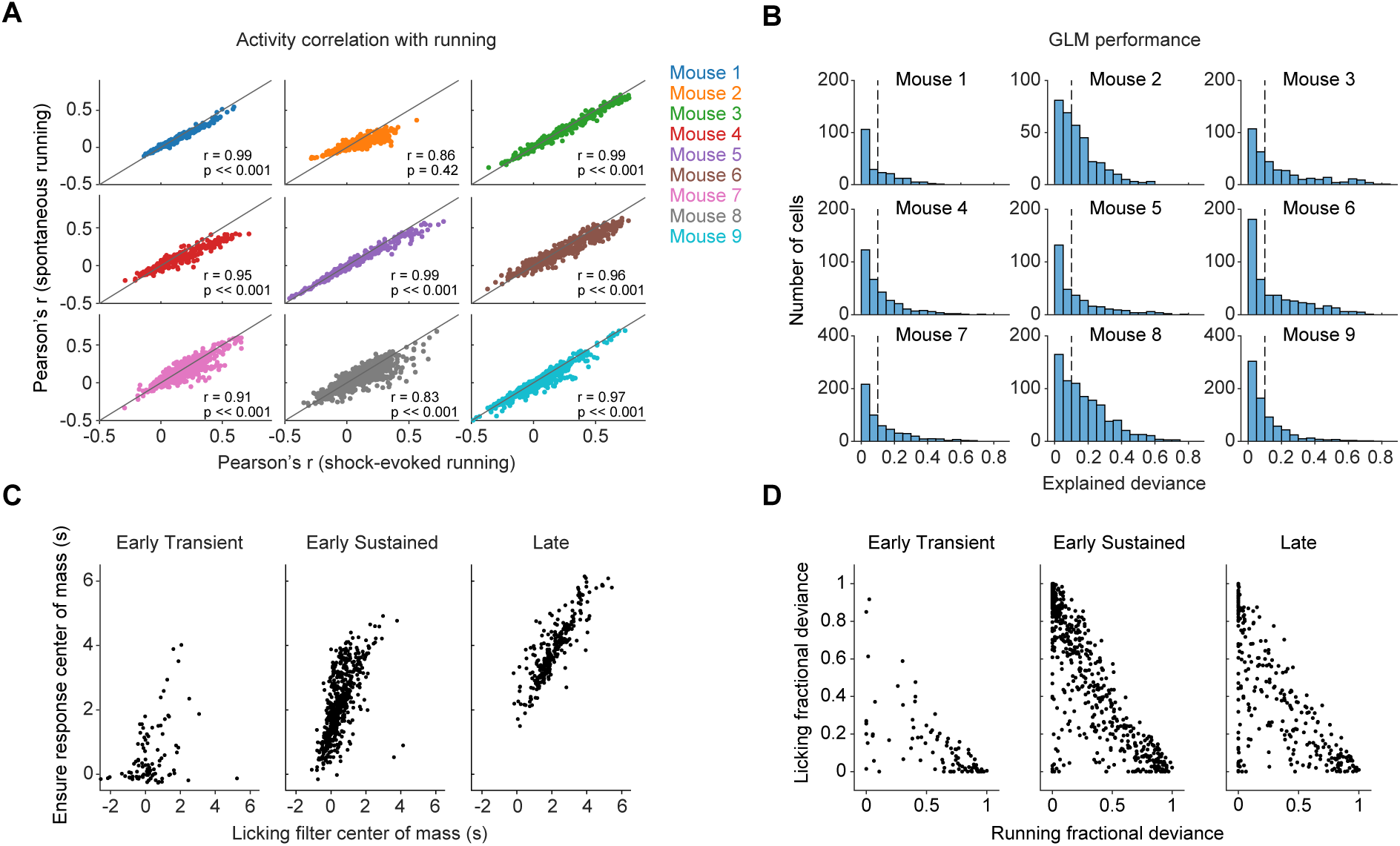
Further analyses of running and licking-evoked activity in LPBN using a GLM. (A) Correspondence of Pearson’s correlation between neural activity and running during the ITI with Pearson’s correlation between activity and shock-evoked running. Each dot is a single neuron. (B) Histogram of GLM explained deviance across neurons for individual mice. Only cells with a model performance of at least 0.1 (dashed line) were used for further analyses. (C) Correspondence of timing in Ensure responses and licking filter delays for each type of Ensure-responsive neuron (n = 1966 neurons from 9 mice). (D) Comparison of fractional deviance for licking versus running for each type of Ensure-responsive neuron (n = 1966 neurons from 9 mice). Fractional deviance is the deviance explained of a model that predicts neural activity with only one behavioral variable (i.e., running or licking) divided by the deviance explained for the full model (including all behavioral variables).

**Figure S5, Related to Figure 5:**
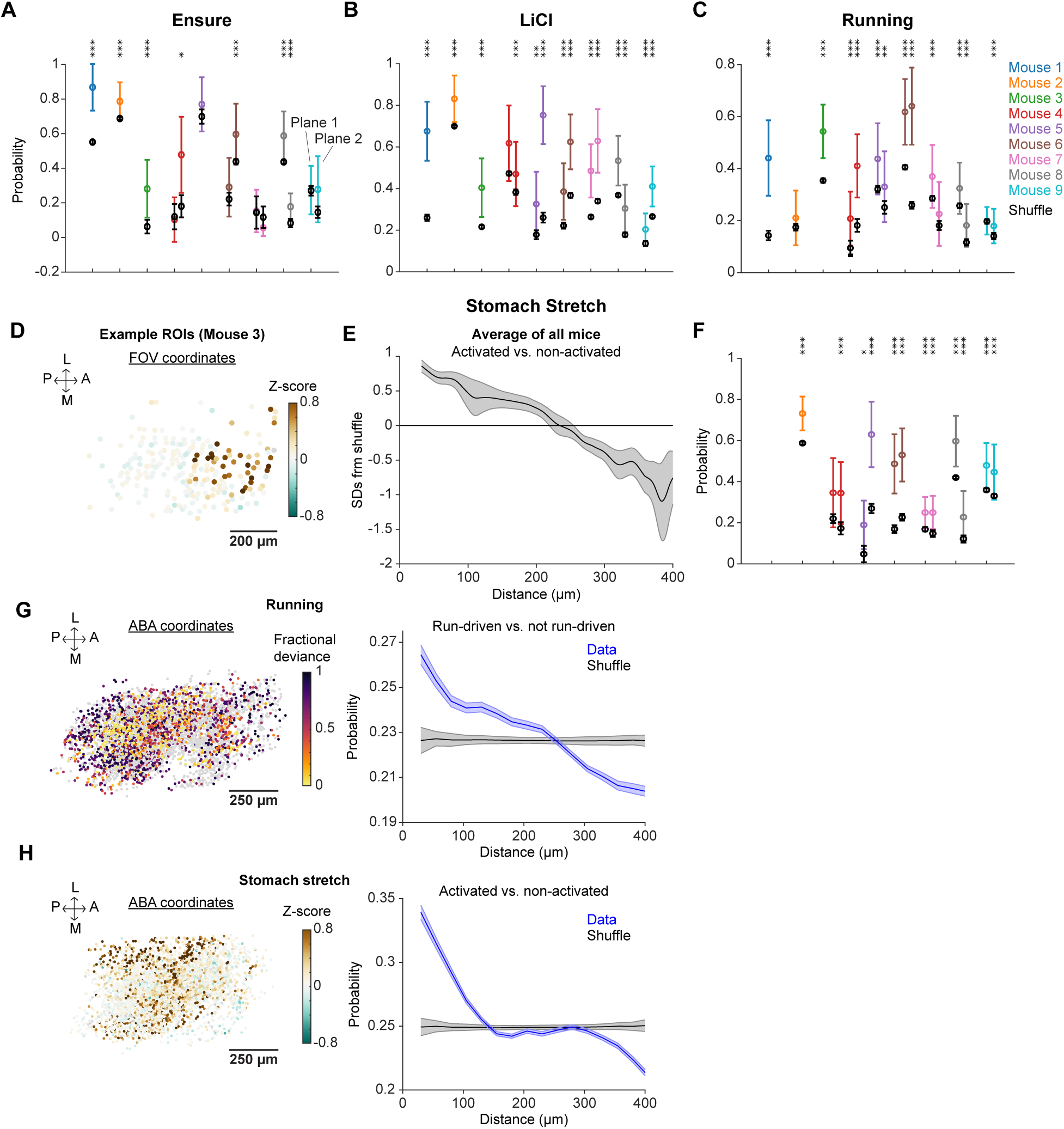
Further analyses of spatial organization of functional subtypes. (A) Likelihood that a pair of neurons spaced <100 µm apart were both Late Ensure (colored dots) compared to shuffled controls (black dots) in FOVs of all mice. We first computed the probability that two neurons within a certain distance of each other were both Late Ensure neurons. Next, we computed the same probability after 100 shuffles of the identity of the neurons. Colored dots show the average probability for all pairs of neurons <100 µm from each other, while black dots show the same for the shuffled data. *P<0.05, **P<0.01, ***P<0.001 comparing data and shuffled data for each mouse with a Wilcoxon rank sum test (see columns of stars at top). P-values were Bonferroni corrected for recordings with two imaging planes. (B) Same as (A) for LiCl-activated cells vs. non-activated cells. (C) Same as (A) for cells driven by running (drivenness defined as a running fractional deviance > 0.65) vs. all other cells. (D) Example cell masks in field of view (FOV) colored by responses to stomach stretch (0-9 s post-inflation onset). (E) Likelihood that two neurons spaced different distances apart were both activated by stomach stretch (expressed as the number of standard deviations [SDs] above shuffle controls), averaged over all mice (n = 7). Shaded regions show SEM across mice. (F) Same as (A) for stomach stretch. (G) Left: Cell masks (n = 3602 from 9 mice) in Allen Brain Atlas coordinates colored by running fractional deviance. Neurons that are not well-explained by the GLM (explained deviance < 0.1) are gray. Right: Likelihood that a pair of neurons spaced different distances apart are both driven by running. Shaded regions show SEM across neurons. (H) Left: Cell masks (n = 3602 from 9 mice) in Allen Brain Atlas coordinates colored by stomach stretch response. Right: Likelihood that a pair of neurons spaced different distances apart are both driven by stomach stretch. Shaded regions show SEM across neurons.

**Figure S6, Related to Figure 6.**
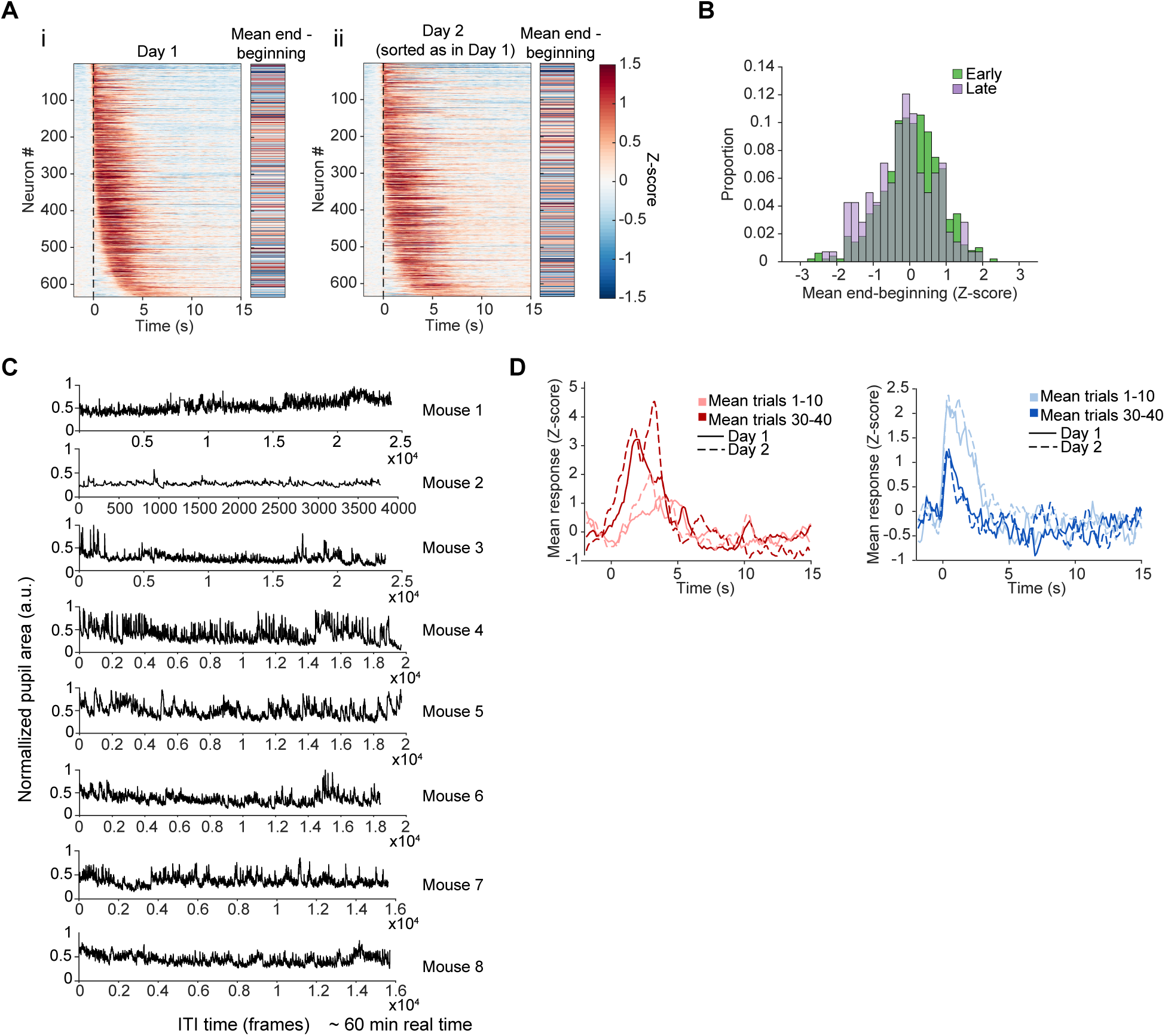
Behavioral controls and further analyses of slow activity changes. (A) Left panel of i: Heatmap showing the mean Ensure response on Day 1 of all Ensure-activated neurons found on two consecutive days of imaging (n = 634 neurons from 8 mice). Right panel of i: Change in mean activity from the last 2 minutes of the session relative to the first 2 minutes of the session of the same neurons. Both panels are sorted by Ensure response onset time. ii: Same as left two panels but for Day 2. Neurons are sorted by Ensure response onset time on Day 1. (B) Distribution of slow activity changes for all Early and Late Ensure neurons. (C) Concatenated pupil area traces over baseline periods and ITI intervals involving quiet waking throughout one session for each mouse. (D) Example neurons that consistently change their Ensure-evoked responses over trials on two consecutive imaging days. Left: Example increase neuron. Right: Example decrease neuron. Lighter lines: Mean response from trials 1-10; Darker lines: Mean response from trials 30-40; Solid lines: Day 1 means; Dashed lines: Day 2 means.

**Figure S7, Related to Figure 7:**
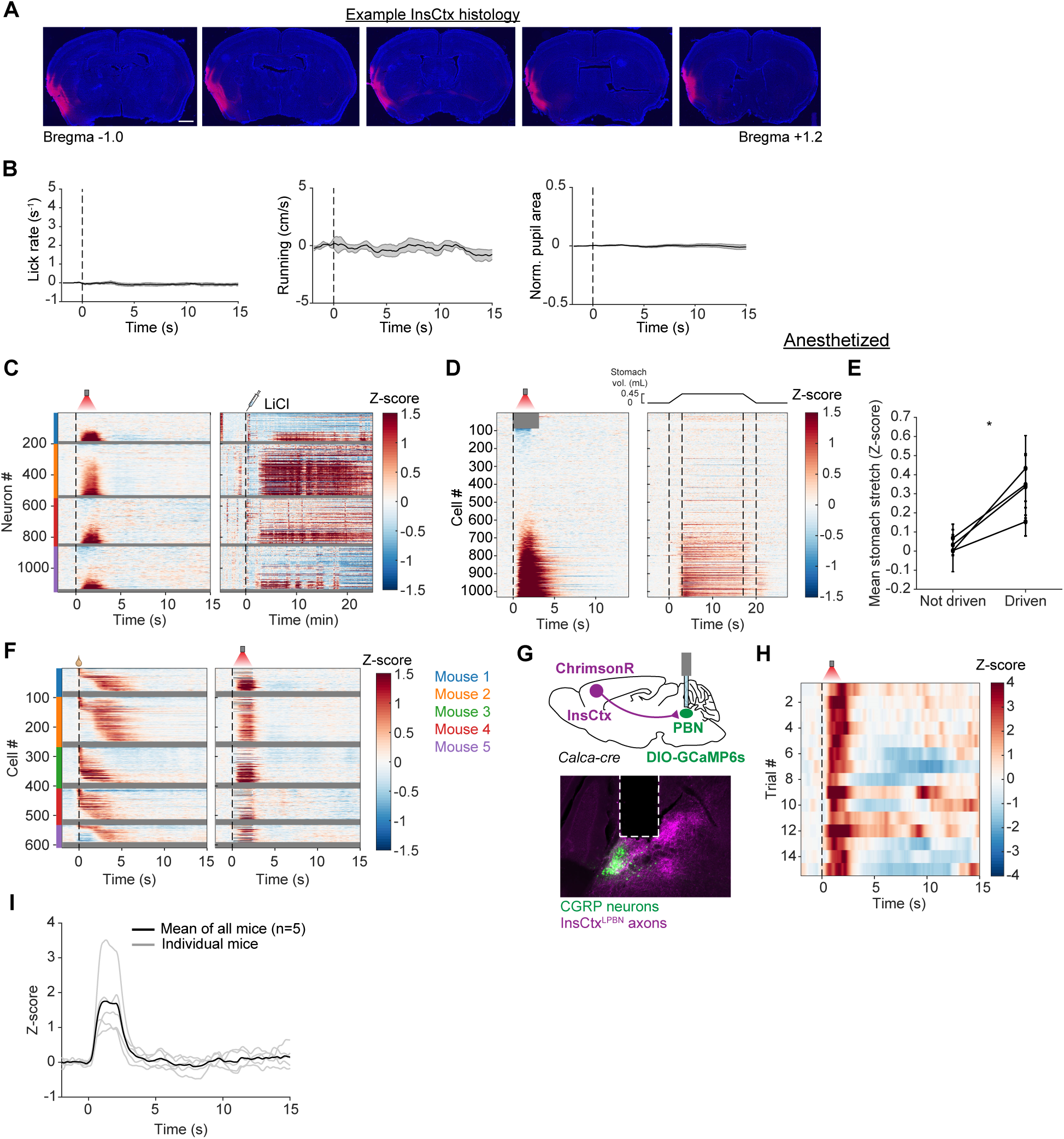
Further analyses of LPBN responses to InsCtx^LPBN^ axon stimulation; CGRP photometry during InsCtx^LPBN^ axon stimulation. (A) Coronal histology showing expression of ChrimsonR-tdTomato throughout the anterior-posterior extent of InsCtx in an example mouse. Scale bar: 250 μm (B) Behavioral responses to InsCtx^LPBN^ axon stimulation. Mean lick rate (left), mean running speed (middle), and mean max-normalized pupil area (right; baseline corrected), aligned to stimulation onset across all trials from all mice (n = 140 total trials from 5 mice). (C) Left: Heatmaps showing the mean response to InsCtx^LPBN^ stimulation of single neurons from individual mice. Right: Heatmaps showing the response of the same neurons to LiCl injection. For both panels, each row is the activity from a single neuron and horizontal gray lines divide the data from each mouse. Neurons are sorted per mouse by the mean Z-score from 0-2 s during photostimulation. Mouse legend is shown in F. (D) Left: Heatmaps showing the mean response of single neurons to InsCtx^LPBN^ stimulation during anesthesia (n = 1030 neurons from 4 mice). Right: Heatmaps showing the response of the same neurons to stomach stretch during the same session. In both heatmaps, neurons are sorted by the mean Z-score from 0-2 s during photostimulation. (E) Mean response to stomach stretch for neurons not driven or driven by InsCtx^LPBN^ stimulation. Each dot is the mean from an individual mouse and values for a given mouse are connected by a line. Dashed line at Z = 0 indicates no response. *P = 0.0148, Paired t test. (F) Left: Heatmaps showing the mean Ensure response of single Ensure-activated neurons from individual mice. Right: Heatmaps showing the response of the same neurons to InsCtx^LPBN^ stimulation. Grayed out responses are from neurons in the region of the FOV where the PMT was blanked during stimulation. For both panels, each row is the activity from a single neuron and horizontal gray lines divide the data from each mouse. Neurons are sorted per mouse by Ensure onset time. (G) Optogenetic stimulation of InsCtx^LPBN^ axons during CGRP neuron fiber photometry recordings. Top: Schematic for ChrimsonR expression in InsCtx and GCaMP6s expression in CGRP neurons with optic fiber implantation above LPBN. Bottom: Coronal histology from an example mouse showing GCaMP6s expression in CGRP neurons, ChrimsonR-tdTomato expression in InsCtx^LPBN^ axons, and optic fiber placement (dashed white lines) above LPBN. (H) Heatmap showing single-trial responses of CGRP neurons to InsCtx^LPBN^ axon stimulation. Each row is the response to a single 2-s stimulation trial. (I) Mean InsCtx^LPBN^ stimulation-induced CGRP neuron activity across all trials from all mice. Gray traces: Mean across trials for a single mouse (n = 15 trials/mouse); Black trace: Mean across all mice (n= 5 mice). Stimulation occurs from 0-2 s. Error bars are +/- SEM unless otherwise indicated.

**Figure S8, Related to Figures 3 and 4:**
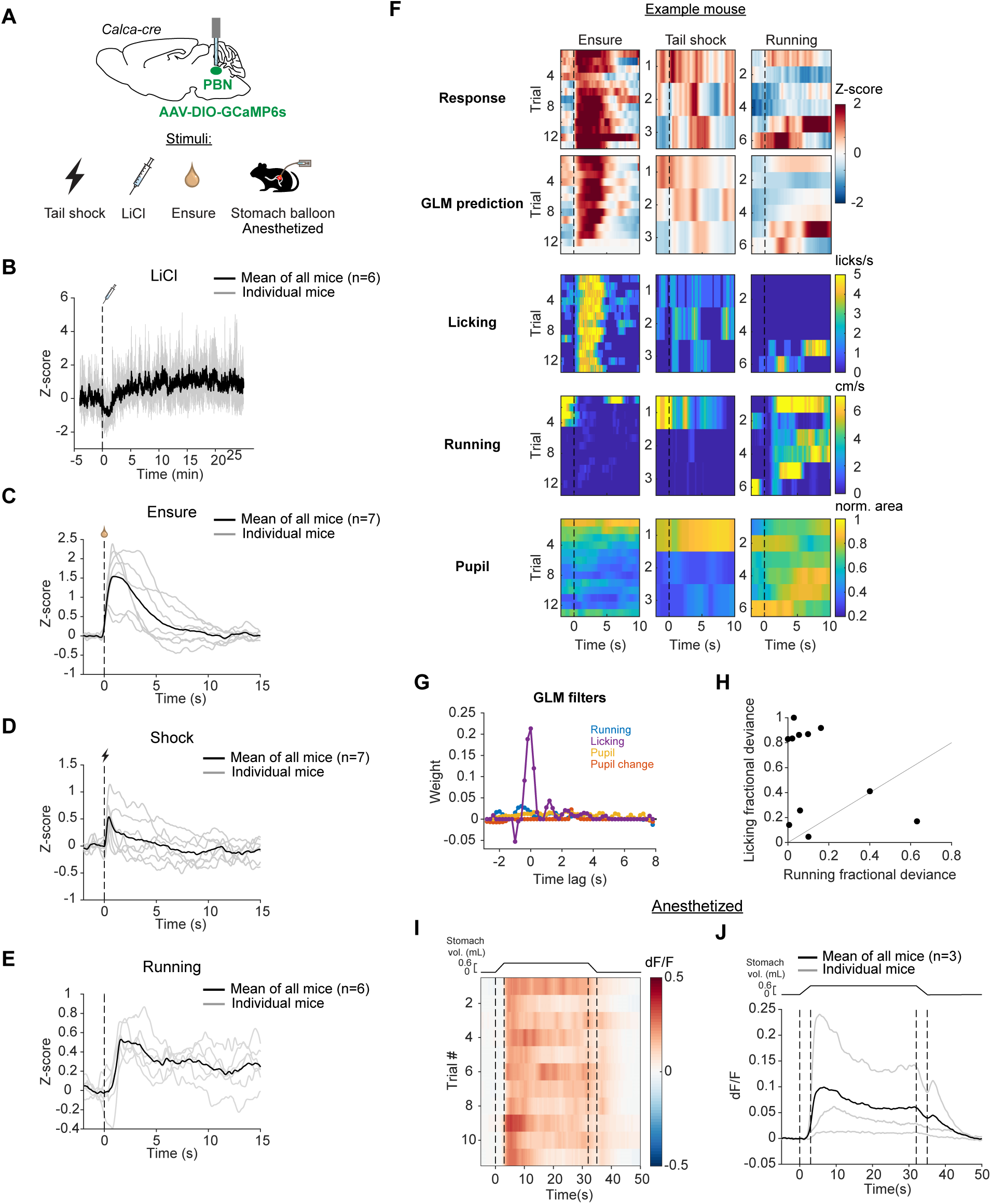
CGRP neuron photometry. (A) Top: Schematic for GCaMP6s expression in CGRP neurons with optic fiber implantation above LPBN. Bottom: Stimuli delivered during photometry sessions. (B) CGRP activity in response to LiCl injection (gray = individual mice; black = average over all 6 mice). (C) CGRP activity in response to Ensure consumption. Gray: mean response across trials (n = 24-40 trials) per mouse. Black: mean response over all 7 mice. (D) CGRP activity in response to tail-shock. Gray: mean response across trials (n = 15-24 trials) per mouse. Black: mean response over all 7 mice. (E) CGRP activity in response to running bouts. Gray: mean response across trials (n = 31-93 trials) per mouse. Black: mean response over 6 mice. (F) Top rows: Heatmaps showing single-trial responses of CGRP neurons to Ensure, tail shock, and running bouts, with corresponding GLM prediction of activity. Bottom rows: Heatmaps showing behavioral predictors (lick rate, running speed, and pupil area (as an index of arousal)) during the stimuli. (G) GLM filters showing the influence of each behavioral variable for the CGRP activity in (F). (H) Comparison of the fractional deviance of licking and running for all photometry mice, showing CGRP activity is more driven by licking than running. (I) Heatmap showing single-trial responses of CGRP neurons to stomach balloon inflation in an anesthetized recording. (J) CGRP activity in response to anesthetized stomach stretch. Gray: mean response across trials (n = 10-14 trials) per mouse. Black: mean response across all mice (n = 3).

## Supplementary video legends

**Video S1. Fluoroscopy during food consumption**

Raw example fluoroscopy recording of an Ensure trial (frame rate: 30 Hz).

**Video S2. Ensure bolus movement from stomach to duodenum**

Movies from 4 mice showing Ensure bolus exit from the stomach into the duodenum on an example Ensure trial. Red arrows indicate moments of duodenum entry. Movies are sped up 2x real-time.

**Video S3. Gradual increase in stomach size during feeding paradigm**

Sped up (10x real-time) fluoroscopy recording during a 15-minute session with 35 trials of Ensure delivery. Note the growing accumulation of food in the forestomach.

**Video S4. Measuring stomach area with DLC.**

Sample fluoroscopy recording with colored points from a custom-trained DLC model tracking stomach edges. Colored lines are spline fits of these points and were used to estimate stomach area, which is shown to gradually increase as the mouse consumes Ensure throughout the session. Movie is sped up 10x real-time.

**Video S5. Ensure-induced dF/F example FOVs.**

Fractional mean change in fluorescence (dF/F) in response to Ensure consumption in FOVs from three mice. Movies show activity from 2 s prior to first lick to 15 s after first lick. Flashing white square indicates time of first lick. Color scale is -0.1 to 0.2 dF/F. Top: lateral; Bottom: medial; Left: posterior; Right: anterior. Each FOV has dimensions ∼800 μm x 500 μm.

